# Heritable variation in root bacterial microbiomes alters natural selection on host plants

**DOI:** 10.64898/2026.10.07.757180

**Authors:** Lauren Carley, Anita Simha, Ethan Bass, Amanda Carmichael, Maggie R. Wagner

## Abstract

Heritable variation in microbiome compositions could theoretically evolve in response to selection acting on hosts, but empirical evidence of this is scarce. We tested for effects of allelic variation in *Boechera stricta* on microbiome composition and downstream consequences for host fitness. In the greenhouse, near-isogenic lines with contrasting phytochemistry alleles differentially conditioned soil microbiota in ways that changed the genotypes’ relative fecundity. In the field, root microbiome differences mediated allelic effects on host fitness. Selection gradients on individual taxa and multivariate axes of root microbiome composition suggest that independent components of the bacterial community favor each allele, especially under drought. These results show that microbiota contribute to natural selection on a major-effect phytochemistry gene, empirically “closing the loop” between microbiome heritability and host trait evolution.

## Introduction

Microbial communities can strongly influence macro-organisms’ physiology and fitness (*1–3*). The evolutionary consequences of this for the host depend on two factors: how much variation in microbiome composition is heritable (controlled by host genes), and how strongly this heritable fraction of the microbiome impacts fitness (Fig. 1A). Although host genetic effects on microbiome composition (hereafter, “microbiome heritability”) are now well documented (*4–7*), their fitness consequences remain largely untested.

**Figure 1.**
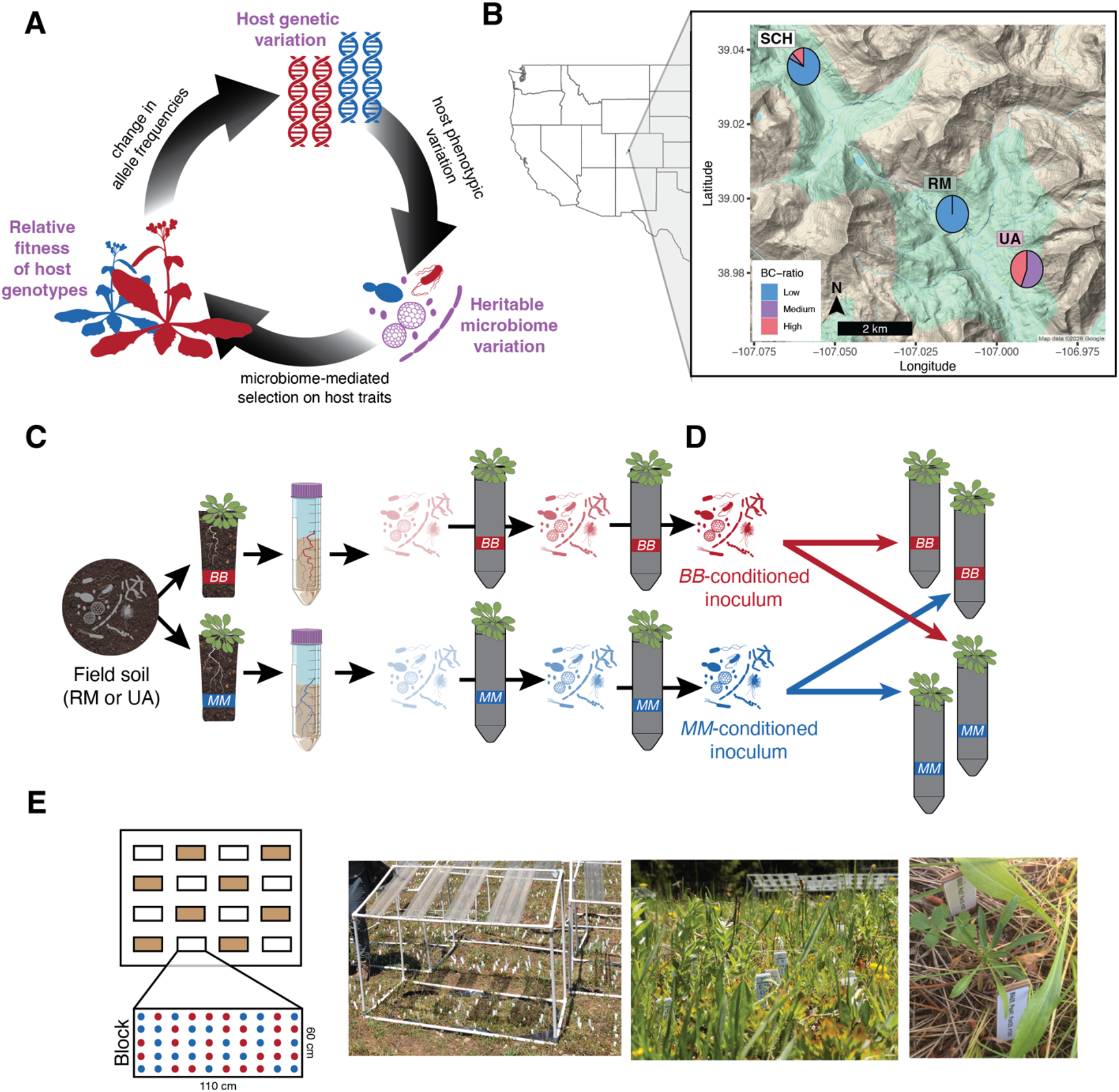
We leveraged an experimentally tractable chemical polymorphism in *Boechera stricta* to test for feedbacks between plant genotype and microbiome composition in greenhouse and field conditions. **(A)** Conceptual figure of project objective. Each arrow represents a biological relationship that has been demonstrated in previous studies; however, studies that “close the loop” by demonstrating that heritable microbiome variation alters the relative fitness of the underlying genetic variants are exceedingly rare. The mechanism for each relationship accompanies each arrow in black text. **(B)** We focus on the role of *BCMA*, a major-effect gene controlling glucosinolate profile quality in *Boechera stricta*. The *BB* and *MM* alleles at this locus are segregating in natural populations in our study region, causing within- and across-population variation in BC-ratio, i.e. the proportion of total aliphatic glucosinolates derived from branched-chain amino acid precursors. Within the study region surrounding the Rocky Mountain Biological Laboratory (Gothic, CO, USA), the three focal field sites are located as points on a map, with inset pie charts showing the local distribution of BC-ratio among wild plants. **(C)** To demonstrate the host genotype → microbiome relationship, we allowed *BB* and *MM* near-isogenic lines to condition two soils collected from natural *B. stricta* habitats and passaged the resulting rhizosphere microbiota forward to inoculate new pots for 3 total cycles of rhizosphere selection. **(D)** The final *BB-* or *MM-*conditioned rhizospheres were aliquotted for microbial characterization and used as inocula in a reciprocal transplant experiment in a growth chamber. We grew both host genotypes in both inoculum types and measured the subsequent expression of several traits and fecundity (flower and fruit production). **(E)** We also quantified the rhizosphere and root microbiomes of *MM* and *BB* near-isogenic lines grown in a common garden in Schofield Meadow, CO, USA, where growth and fecundity were recorded over 4 years (*31*). *From left to right:* Schematic of the common garden experiment, with *MM* and *BB* genotypes (red and blue dots) grown in a randomized complete block design, with half the blocks receiving a precipitation-reduction treatment (rainout shelters, brown boxes); Photograph of rainout shelters over transplanted *BCMA* NILs; Photograph of experimental plants early in the growing season, surrounded by natural neighboring vegetation, with rainout shelters visible in the background; Photograph of a single rosette. In all three photographs, plastic tags with white labels mark each transplanted individual. Vector art of organisms courtesy of Arcadia Science and PhyloPic.

Several studies have generated correlational evidence consistent with the hypothesis that microbiome variation may adaptively evolve *via* selection on the host (*8*, *9*). However, direct empirical tests remain rare (*10*) because the experiments required to prove that heritable microbiome differences causally shape the relative fitnesses of host genotypes are challenging (*11*). A few notable studies have taken the key step of demonstrating that experimental microbiome manipulation can influence host fitness or evolution. For example, coevolution has been experimentally demonstrated in the highly specialized legume-rhizobia symbiosis (*12*, *13*). In *Drosophila melanogaster*, different bacterial inoculants drove genome-wide allele frequencies in opposite directions (*14*). A recent field study in *Arabidopsis thaliana* found that heritable variation for foliar bacteria contributes substantially to genetic variance in fecundity (*15*). However, all of these landmark studies emphasize the contribution of particular microbial taxa, and it remains unclear how well these cases represent the heritability and adaptive value of full microbiomes. In contrast, transplanting whole *Daphnia* gut microbiomes between genotypes influenced survival and reproduction in the lab (*16*), but the importance of such effects in natural environments remains unknown. Finally, the emergent functions of heritable whole-microbiome variation have been tested using the intraspecific plant-soil feedback (PSF) framework (*17*, *18*). In PSF studies it is uncommon to directly measure either the conditioned microbiomes or the causal plant genes or traits, but evolutionary consequences of differential conditioning can be quantified through measurement of plant fitness (*19*).

Here, we combine elements from these previous studies to empirically close the loop between microbiome heritability and host fitness (Fig. 1A) and explore whether microbiome composition can evolve as an “extended phenotype” *via* selection acting on the host. In the perennial wildflower *Boechera stricta* (Graham) Al-Shehbaz (*35*), we tested the effects of glucosinolate chemistry on root-associated microbiomes and the resulting feedbacks on host fecundity in the greenhouse and field. Glucosinolates are evolutionarily important phytochemicals that mediate interactions of brassicaceous plants with herbivores, pathogens, and non-pathogenic microbes (*20–25*). Different glucosinolates vary in bioactivity (*26*) and elicit wide-ranging responses from microbes (*27–29*), creating potential for a diversity of glucosinolate-driven plant-microbiome interactions in complex natural communities. In *B. stricta*, natural variation at a large-effect locus, *BCMA1/3* (hereafter, *BCMA*), is maintained by balancing selection throughout the species range (*30*, *31*). *BCMA* controls a major axis of variation in glucosinolate profile: “BC-ratio”, i.e., the proportion of aliphatic glucosinolates that are derived from branched-chain amino acids instead of methionine (*32*, *33*). BC-ratio is polymorphic both within and across *B. stricta* populations, including in the region of this study (Fig. 1B). Balancing selection on *BCMA* is shaped by both herbivore pressure and drought stress; *M* and *B* alleles—producing methionine-derived and branched-chain glucosinolates respectively—each confer stronger herbivore resistance in some environments, while *M* alleles improve fitness under drought (*30*, *31*). The mechanism by which BC-ratio alters water relations is unknown; root-associated microbes can influence plant performance under drought (*34*, *35*), but the potential contribution of the root microbiome to drought response and resulting abiotic selection on *BCMA* has not yet been investigated.

Leveraging *BCMA* near-isogenic lines (NILs), we used a greenhouse experiment to ask whether rhizosphere microbial communities conditioned by contrasting NILs affect plant phenotypic expression and the relative fitness of causal *BCMA* alleles. We then grew the same NILs for four years in a common garden to assess differential microbiome conditioning and its relationship to lifetime fecundity in the field. Because *B. stricta* is predominantly self-pollinating, heterozygosity is naturally low (*36*, *37*) and fitness differences between homozygous NILs reasonably approximate selection in wild populations. By manipulating drought stress using a rainfall exclusion treatment, we also tested how the abiotic environmental context influences the relationships between host-filtered microbiome composition and natural selection (*38*). Finally, by estimating selection gradients on heritable microbiome components and individual microbial taxa (*39*, *40*), we formalized the observed eco-evolutionary feedback between root-associated bacteria and host phytochemistry in this system.

## Results

### *BCMA* controls root glucosinolate profiles

The role of *BCMA* in controlling root chemistry has not yet been explored, as prior molecular and evolutionary characterization of this gene has focused on leaves (*32*, *33*). Paired leaf and root samples from NILs homozygous for alternative *BCMA* alleles (*BB* or *MM*) showed that BC-ratio is strongly correlated between leaves and roots and that root glucosinolate profiles diverge between *MM* and *BB* genotypes (fig. S1) as has previously been described in leaves. Glucosinolates are concentrated in the peripheral cell layers of secondary roots, which senesce during normal development and release glucosinolates into the rhizosphere where they are hydrolyzed by plant and/or microbial myrosinases (*41–43*). Therefore, variation in root glucosinolates likely determines concentrations of glucosinolates and associated breakdown products that microbes encounter in rhizosphere soil.

### *BCMA*-driven rhizosphere conditioning alters leaf number and fecundity in controlled conditions

To test whether *BCMA* alters the rhizosphere microbiome, we grew *MM* and *BB* NILs in autoclaved potting soil inoculated with field soils from two sites in the Rocky Mountains where local *B. stricta* populations are polymorphic for BC-ratio, yielding two independent microbial pools (Fig. 1B; “UA” and “RM”). We conditioned the starting microbial pools on each host genotype for three consecutive cycles, using the resulting rhizosphere communities to inoculate a new plant of the same genotype (Fig. 1C). Rhizosphere bacterial community composition shifted in response to the conditioning plant’s genotype (Fig. 2); fungal communities showed a qualitatively similar response but with weaker statistical support (fig. S2; table S1). Overall, 88 bacterial ASVs were sensitive to host *BCMA* genotype (Fig. 2C; table S2), comprising 7.9% and 7.1% of the rhizosphere communities derived from each of the two soils. Notably, the effect of *BCMA* genotype on bacterial community composition depended on the soil origin of the microbial inoculum (Fig. 2B), with 55 ASVs showing soil-specific responses to host genotype (Fig. 2D). Some of these ASVs were enriched by *BB* plants in one microbial pool but enriched by *MM* plants in the other, while other ASVs responded to host genotype in only one microbial pool (Fig. 2D). These genotype-environment interactions mirror past findings (*44*, *45*) highlighting how the initial species pool modulates effects of host genotype on microbiome composition.

**Figure 2.**
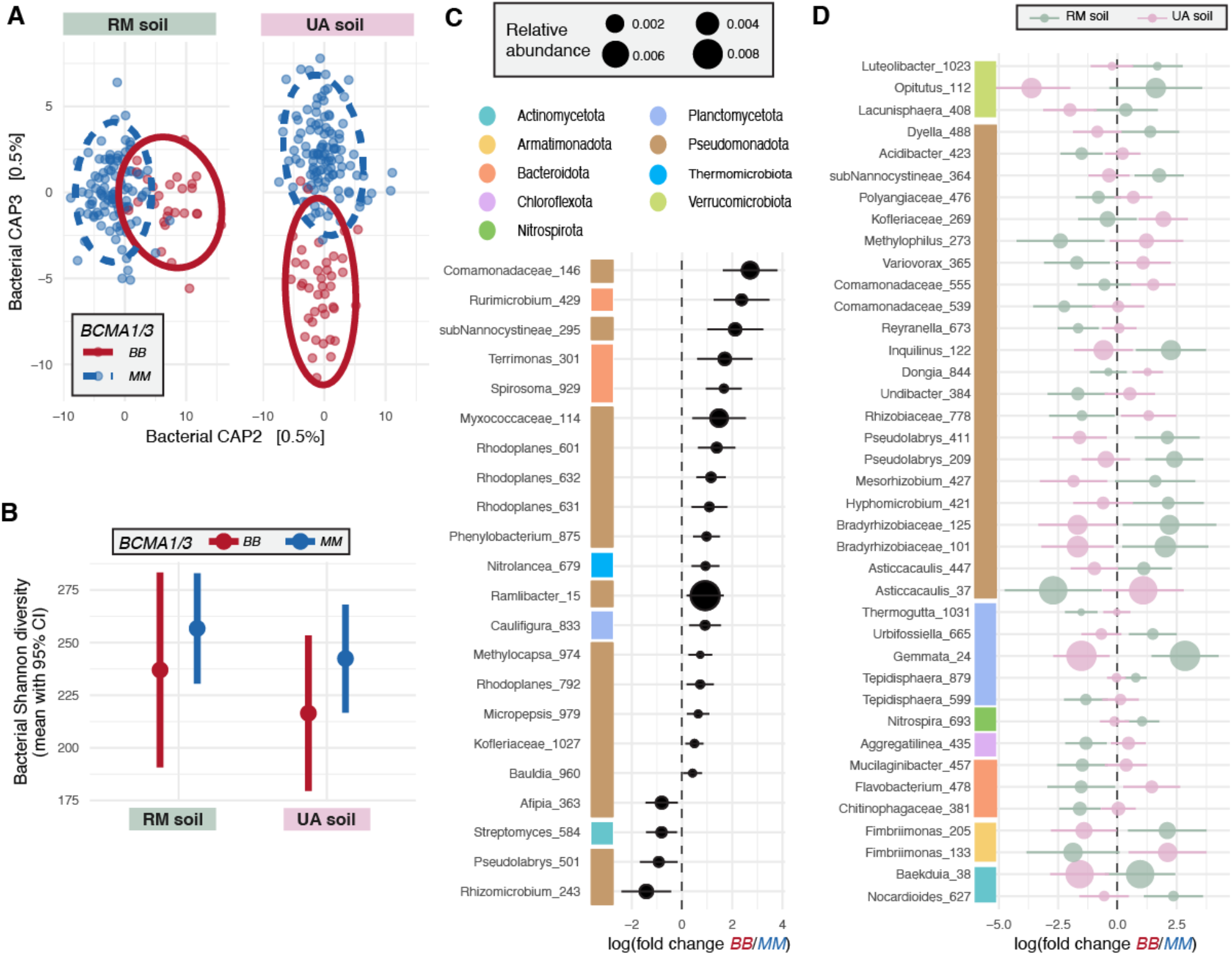
Rhizosphere bacterial communities are differentially conditioned by near-isogenic lines with contrasting root glucosinolate profiles. **(A)** Constrained partial ordination of Aitchison distances (with 95% confidence ellipses) among bacterial rhizosphere communities of *BCMA* NILs. Community composition shifted in response to conditioning genotype (redundancy analysis, *F*_1,276_ = 1.44, P=0.008) and to the conditioning genotype’s interaction with the starting soil pool (redundancy analysis, *BCMA\**Soil interaction *F*_1,276_ = 1.35, P=0.01). Effects of sequencing depth and passaging cycle were partialled out before constraining by genotype, soil, and their interaction. **(B)** Estimated marginal mean alpha diversity (*e*^Shannon^ ^index^) of rhizosphere bacterial communities derived from two field soils and conditioned by *BCMA* NILs. Statistical support is provided in table S1. **(C-D)** Estimated marginal mean changes in bacterial ASV abundance between rhizosphere soils conditioned by *BB* and *MM* genotypes, separated by (C) taxa with a consistent response to *BCMA* across soils and cycles, and (D) taxa with a significant *BCMA\**soil interaction across cycles, grouped by phylum (linear mixed models, *P*_FDR_<0.05). ASVs with *BCMA\**cycle interactions are not shown but are listed in table S2.

To investigate whether the *BCMA*-driven changes in the bacterial microbiota feed back to shape relative fitness of the causal alleles, we conducted a reciprocal re-inoculation experiment in the growth chamber (Fig. 1D). Microbial communities that had been conditioned by either *BB* or *MM* plants (the “conditioning genotype”) were re-inoculated into pots containing either *BB* or *MM* seeds (the “test genotype”), which we tracked throughout the life cycle. Juvenile size, height at flowering, and flowering time were all insensitive to *BB*-conditioned versus *MM*-conditioned microbial treatments (table S3). Adult leaf number in the test generation varied significantly between *BB-*conditioned and *MM-*conditioned microbiota, although effects differed qualitatively between microbial pools (Fig. 3A). This suggests that leaf number, a functional trait that predicts water use under drought (*30*), plastically responds to soil microbiota, including shifts in bacterial composition that are both genotype- and soil-specific.

**Figure 3.**
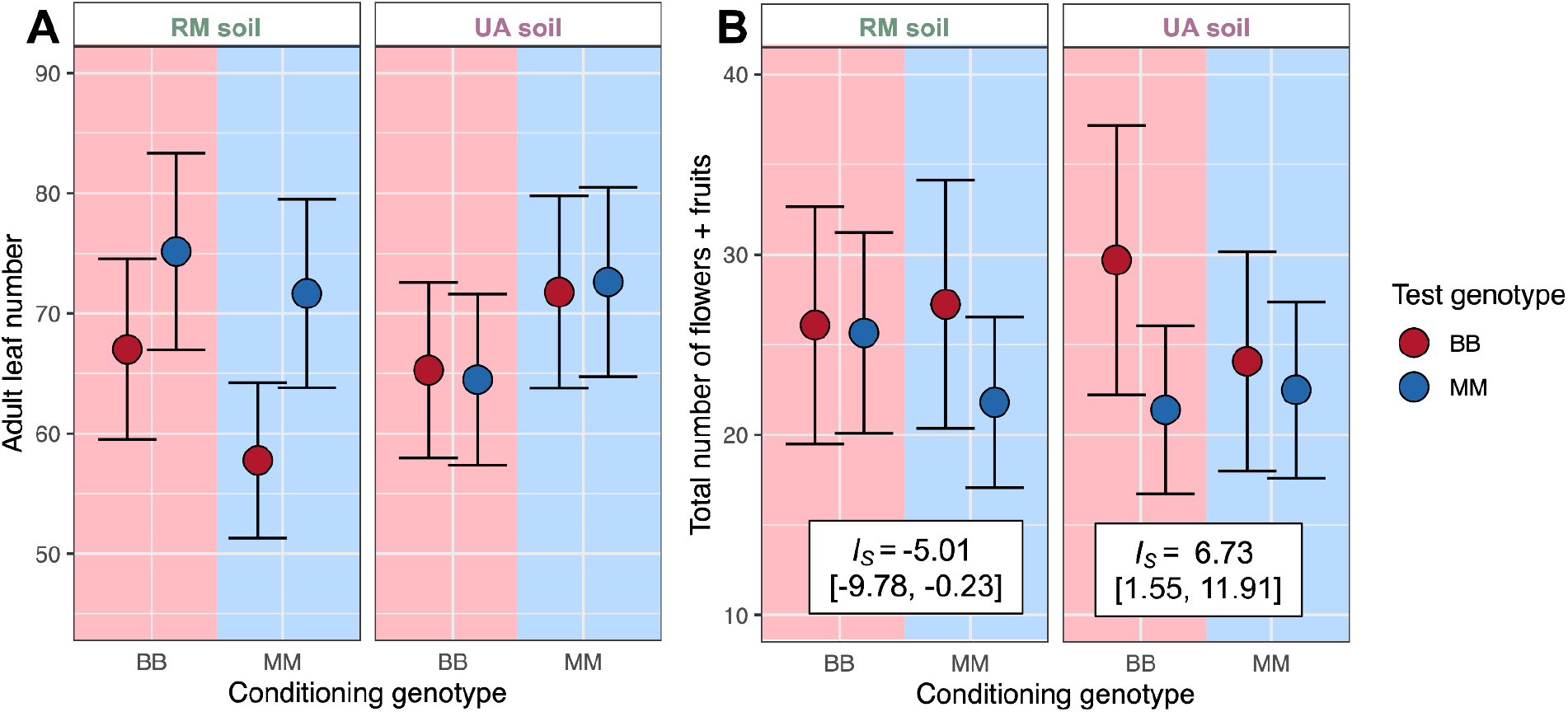
Host-conditioned bacterial microbiomes shape genetic variation for host phenotypes and fitness. **(A)** Leaf number, a trait predicting water use under drought (*36*), was plastic in response to the soil microbiome, including microbiome differences attributable to both the starting soil pool and the conditioning genotype (Genotype_cond_*Soil: *χ*^2^_1_=13.40, *P*=0.0003). In the RM microbial pool (left), test genotypes differ in leaf number, but the absolute number of leaves and strength of genetic differences in leaf number are both increased by inoculation with *BB*-conditioned microbiomes. In the UA microbial pool (right), *MM* but not *BB* test plants showed microbiome-induced plasticity for leaf number, producing significantly more leaves when inoculated with *MM-*conditioned microbiomes. **(B)** Genetic variation for fecundity (total number of fruits and flowers) varied significantly in response to interactions between the conditioning genotype, the test genotype, and the starting microbial pool (Genotype_Test_*Genotype_Cond_.*Soil interaction, *χ*^2^_1_=13.84, *P*=0.0002). Full details on statistical support is provided in table S3. In all panes, points indicate estimated marginal means and bars are +/-1 SE. Text in white boxes in **B** are quantitative interaction coefficients (point estimate and 95% confidence intervals) parameterizing the plant-soil feedback in each microbial pool (Bever *et al.* 1997).

The total number of reproductive structures was shaped by three-way interactions between conditioning genotype, test genotype, and starting microbial pool (Fig. 3B), implying that genetic variation for host fecundity depends on the interaction between conditioning genotype and the starting microbial pool. Interestingly, the evolutionary implications of this feedback differed between the two starting microbial pools. In the UA microbial pool, the *BB* genotype conditioned the rhizosphere in a way that increased its own relative fecundity (*I_S_*=6.73: a positive feedback, which would promote fixation of *B* alleles (*19*)). In the RM microbial pool, the *MM* genotype conditioned the rhizosphere in a way that reduced its own fecundity relative to *BB* (*I_S_*=-5.01: a negative feedback, which could maintain allelic diversity (*19*)). Together, these results demonstrate that genotype-specific microbiome conditioning can significantly alter natural selection on the causal genotypes, resulting in either positive or negative feedbacks depending on the initial microbial community.

### Root endosphere bacteria mediate *BCMA* effects on host fecundity in the field, especially under elevated drought stress

To determine whether the observed *BCMA*-driven rhizosphere conditioning effect also occurs in natural environments, we grew the same NILs in a common garden in Schofield Meadow, CO (Fig. 1E), where the local population is polymorphic for BC-ratio (Fig. 1B). To investigate interactions between *BCMA* genotype and abiotic stress, half of the randomized experimental blocks received partial rainfall exclusion, which reduced soil moisture (*30*). We measured fecundity (fruit set) annually and then censused rhizosphere and root endosphere microbiomes of 90 surviving individuals at the end of the 4-year-long experiment. As in the growth chamber experiment, *BCMA* influenced the taxonomic composition of rhizosphere bacterial communities in the field (table S4; fig. S3), confirming that microbiome conditioning can happen within one generation. Fungal community composition also differed between *BB* and *MM* rhizospheres in the field (table s4; fig. S4), unlike in the growth chamber (table S1B). Host *BCMA* genotype also shaped the bacterial and fungal root endosphere microbiomes (figs. S3, S4), as would be expected if glucosinolates are exuded from the root into the rhizosphere (*46*) from which the endosphere community is derived (*47*).

Consistent with past observations (*30*, *31*), the relative lifetime fecundity of *BB* and *MM* NILs reversed depending on water availability, with *BB* favored under ambient field conditions and *MM* favored under elevated drought stress (Fig. 4A). Drought treatment also reversed the direction of host genotype-driven shifts in microbiome composition (Fig. 4B). Motivated by the results of our microbiome reciprocal transplant experiment (Fig. 2F), we used structural equation modeling to separate the direct effect of *BCMA* on relative fecundity from indirect effects mediated through the bacterial microbiome (quantified as canonical analysis of principal coordinates [CAP] axes; table S4. Both CAP3 and CAP4 of the bacterial root endosphere microbiome were associated with host fecundity in the drought treatment (Fig. 4C-D; table S5), indicating directional selection on these components of the microbiome (*40*, *48*).

**Figure 4.**
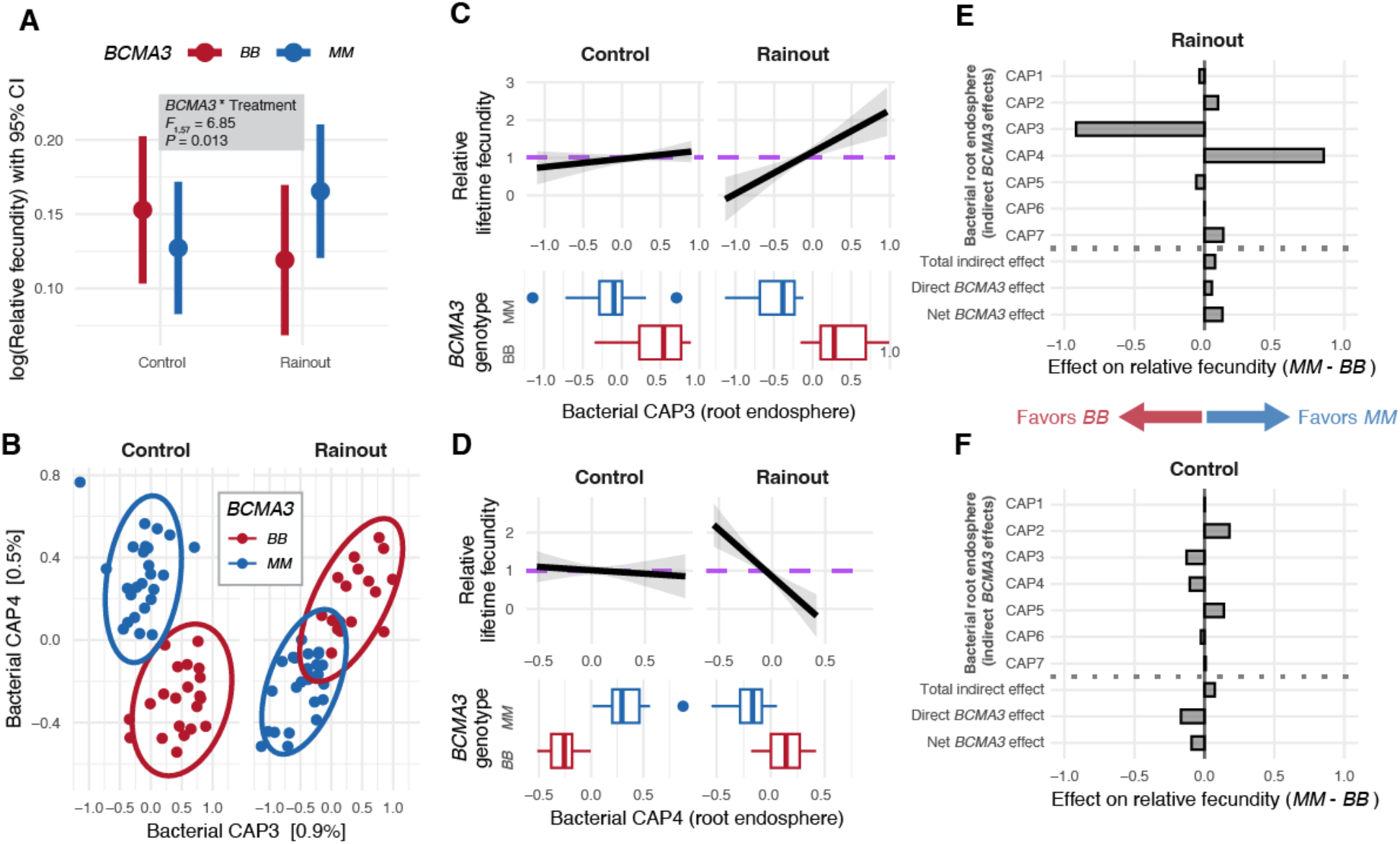
*BCMA*-driven variation in root endosphere bacterial microbiomes mediated lifetime fecundity in field-grown plants under reduced precipitation. **(A)** Supporting previous work, the relative lifetime fecundity of *BB* versus *MM* plants reversed between control and rainout treatments, with the *MM* genotype enhancing fitness under reduced precipitation (rainout) conditions. Points show estimated marginal means with 95% confidence intervals from a model that used no information about microbiome composition. **(B)** In both control plots and rainout plots, the third and fourth constrained axes of variation (CAP3 and CAP4) differentiated the root endosphere microbiomes of *BB* and *MM* plants. For CAP4, the *BCMA* effect reversed direction between treatments. Detailed results from the canonical analysis of principal coordinates are provided in table S4 and fig. S3. **(C-D)** Both CAP3 and CAP4 predicted relative lifetime fecundity of host plants in the rainout treatment. The purple dashed line represents the null hypothesis of no association between relative fecundity and microbiome variation. Selection gradients are drawn as black lines, with the standard error in grey. The boxplots near the horizontal axes show the distributions of CAP3 and CAP4 axis scores for host plants with contrasting *BCMA* genotypes. Statistical support and selection gradient estimates for other microbiome axes are provided in table S5. **(E-F)** Structural equation modeling identified that both CAP3 and CAP4 mediated the effect of *BCMA* on relative lifetime fecundity in the rainout treatment (E; CAP3: Δ⍵ = -0.919 [95% CI -2.548, -0.038]; CAP4: Δ⍵ = 0.964 [95% CI 0.255, 2.083]), but in opposite directions. In contrast, we detected no evidence for microbial mediation in the control treatment (F). Statistical support is provided in table S6.

The multiplication of these selection gradients by the effects of host genotype on the corresponding axes (Fig. 4B) revealed strong indirect effects of *BCMA* on relative fecundity *via* the gene’s influence on these two axes of the root bacterial microbiome (Fig. 4E; table S6). Notably, the two CAP axes had opposite effects on the relative fitness of the *BCMA* alleles such that the total indirect effect of *BCMA-*driven microbiome variation was close to zero. This indicates that under water-limited conditions, two independent components of the root bacterial microbiome respond to glucosinolates and impact host fecundity, but one (summarized by CAP3) favors *BB* genotypes and the other (summarized by CAP4) favors *MM* genotypes. By contrast, we found no evidence that root bacterial microbiome variation mediated fitness effects of *BCMA* in the control treatment (Fig. 4F; table S6). This observation supports findings that bacterial impacts on plant phenotype and fitness are stronger under drought stress (*34*) and may reflect legacy effects of water availability on soil microbiome function (*49*) and/or altered physiological interactions (*50*). Together, these results suggest that heritable components of the *B. stricta* root bacterial microbiome may contribute to balancing selection on *BCMA* under drought stress.

### Selection on bacterial endosphere ASVs is taxonomically and environmentally patterned

To investigate which bacterial taxa mediate the fitness effects of *BCMA* under reduced precipitation (Fig. 4C-E), we back-projected the selection gradients (*β*) onto the CAP axes to estimate selection gradients on the constituent ASVs (*39*) in the root endosphere under drought. We found strong evidence for negative selection on the relative abundances of 156 bacterial ASVs and positive selection on 133 ASVs (table S7). These conservatively-defined sets comprised 9.1% and 7.7% of the 1,719 bacterial ASVs, respectively. At higher levels of taxonomic organization, more groups had positive mean selection gradients than negative (Fig. 5A). However, for most groups the mean selection gradient was near zero, indicating either that ASVs in these taxa did not contribute to host fitness or that the taxon contained a mix of positively- and negatively-selected ASVs.

**Figure 5.**
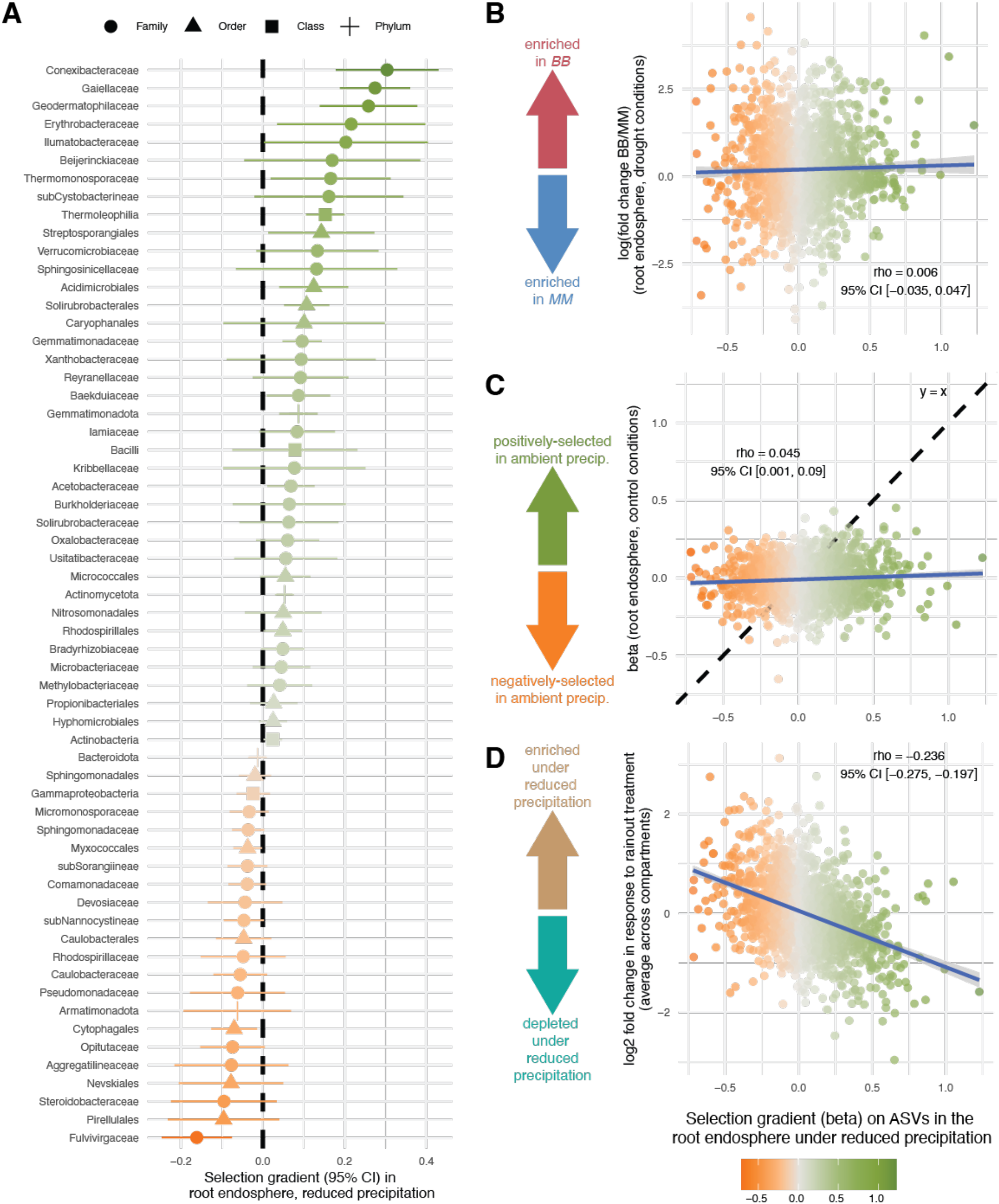
Selection gradients on bacterial ASV relative abundances in the root endospheres of droughted plants are taxonomically and environmentally structured. **(A)** The estimated marginal mean selection gradients on bacterial root endosphere ASVs within 60 higher-level taxa (taxonomic level indicated by shape) are shown. Mean selection gradients differed among bacterial orders (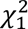=12.31, *P*=0.0004), classes (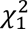=6.13, *P*=0.013), and phyla (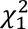, *P*=6.71e^-6^). Taxa with fewer than 5 representative ASVs in our dataset were excluded from the analysis, and 40 taxa with mean selection gradients close to zero and/or with weak statistical support are omitted for figure clarity; however, results for all bacterial ASVs are available in table S7. **(B-D)** Spearman correlations of selection gradients on 1,719 bacterial ASVs in the root endosphere of droughted plants (horizontal axis) with **(B)** the difference in ASV relative abundance between host genotypes; **(C)** selection gradients on ASVs under control rather than dry conditions; and **(D)** the change in ASV relative abundance due to experimental drought. Estimates and confidence intervals for *ρ* are based on 10,000 Monte Carlo draws, in which *x* and *y* values for each ASV were resampled from normal distributions centered on each point estimate with standard deviation equal to its associated standard error, and are likely attenuated toward zero.

Enrichment of negatively- and positively-selected ASVs was balanced between host genotypes (Fig. 5B), supporting our previous observation that separate components of the root microbiota positively mediate fitness effects of each allele (Fig. 4E-F). Endophyte associations with host fitness strongly depended on water availability. ASV selection gradients had greater magnitude under water-limited conditions (Fig. 5C), consistent with evidence that drought can strengthen the impacts of both beneficial and pathogenic bacteria (*34*). Surprisingly, drought enriched negatively-selected ASVs while depleting positively-selected ASVs (Fig. 5D). This suggests that water limitation may weaken host control of the microbiome by suppressing plant defenses (*51*) and/or inducing virulence in opportunistic endophytes (*52*), challenging alternative hypotheses predicting selective recruitment of mutualists to mitigate stressors (*53*). In this study, experimental drought also worsened herbivory damage (*31*) but it is unknown whether this is due to suppression of glucosinolates—which could also reduce defense against weakly pathogenic microbes. In general, evidence remains mixed whether and when drought stress induces or suppresses the expression of plant chemical defenses.

## Discussion

Observations of heritable microbiome variation are common, but cases where such variation clearly affects host fitness are still rare. Here, we showed that (1) contrasting alleles of a major-effect phytochemistry gene in the wildflower *Boechera stricta* differentially shape the composition of the root bacterial microbiome (Figures 2, 4), and (2) these changes feed back to affect the relative fitness of the two causal alleles (Figures 3, 4). Together these results fulfill the necessary conditions for evolution of plant-microbe associations as “extended phenotypes” *via* natural selection acting on the host plant (*54*). They also constitute a rare example of an eco-evolutionary feedback (EEFs) altering natural selection in the wild (*55*). Strikingly, both the allele-specific bacterial conditioning (“evo-to-eco”) and microbiome effects on genetic variation for fitness (“eco-to-evo”) observed in growth chambers were recapitulated under natural field conditions.

Our results provide strong evidence that EEFs may drive coupled evolution of phytochemistry and the microbiome “extended phenotype” in plants. Such feedbacks can either maintain allelic variation via negative feedback loops (when allele-specific microbiome responses favor the alternative allele) or reduce variation via positive feedback loops (when allele-specific microbiome responses favor the same allele) (*19*). We observed both types of feedback in our reciprocal microbiome transplant experiment, where the outcome depended on the starting microbial community pool (Fig. 3B). This result suggests two mechanisms through which soil microbiome variation could contribute to maintenance of the *BCMA* polymorphism at different spatial scales. Among populations, positive feedbacks could promote local fixation of either allele (directional selection) depending on the ambient microbial pools, potentially maintaining both alleles across a microbially heterogeneous landscape (*56*). Within populations, negative feedbacks (*e.g*., as elicited by *MM* conditioning in RM soil; Fig. 3B) could contribute to local maintenance of *BCMA* polymorphism (balancing selection). Indeed, our field experiment showed that even within a single soil background, each allele can simultaneously be associated with both positive and negative effects on host fitness that are mediated by orthogonal axes of microbial community composition. This suggests that different subsets of co-occurring microbial taxa could mediate parallel EEFs with different evolutionary implications. Future research exploring the spatiotemporal dynamics of microevolutionary change and microbial associations in wild plant populations will help to clarify how microbial feedbacks modify selection and ultimately shape host genetic variation.

These feedbacks between microbiome conditioning and selection on host chemical polymorphism are also shaped by abiotic variation. Building upon previous work showing that balancing selection on *BCMA* is shaped by interactions between herbivory and drought stress (*35,36*), our field study found heightened effects of microbiome conditioning on plant fitness under experimental drought (Fig. 4E-F). Thus, the previously-discussed contributions of microbial feedbacks to maintenance of *BCMA* polymorphism may be strongest in environments with lower water availability and/or higher drought frequency, and in particularly dry years. These effects may be partially mediated by morphological plasticity—as seen in our greenhouse experiment, where *BCMA*-mediated microbiome conditioning was associated with reduced leaf number (Fig. 3A), a trait closely tied to water use in this species (*30*). Glucosinolates may thus influence drought tolerance through multiple pathways: differential conditioning of root-associated microbiomes as well as intrinsic effects on physiology (*e.g*., regulation of stomatal behavior in leaves (*57–59*) and hydraulic conductivity in roots (*60*, *61*)). Future work quantifying water relations in different microbial contexts will clarify the relative contributions of glucosinolates’ direct vs. indirect effects to drought tolerance.

Our results suggest several exciting avenues for further exploration of host-microbiome evolutionary dynamics. First, although we estimated fitness exclusively as fecundity, microbes are expected to influence host survival as well. Future research leveraging experimental designs that incorporate both viability and fecundity selection will improve our understanding of host-microbe EEFs. Second, because this study focused on a single trait (BC-ratio), our findings likely capture only a fraction of the phenotypic diversity shaping microbiome composition. Thus, the total fitness consequences of heritable microbiome variation likely exceed those described here. Finally, while our microbiome transplant experiment in the greenhouse confirmed a causal relationship between microbiome conditioning and fecundity, follow-up work with more diverse genotypes can disentangle genotypic selection patterns from environmental covariances between fecundity, microhabitat properties, and microbial communities in the field (*62*). Future directions such as these will build upon our findings, which provide strong foundational evidence that pleiotropic effects on belowground microbial communities influence the evolution of a major phytochemistry locus in a wild plant, empirically closing the loop between microbiome heritability and host fitness.

## Acknowledgements

We are grateful to Tom Mitchell-Olds for early guidance, support, and input on the project; to Emily A. Beck, Jim D. Bever, John K. Kelly, Rebecca Leuschen-Kohl, and Joel F. Swift for helpful feedback on the manuscript; and to Leana Amezquita, Natalie E. Ford, Ali Gomez Zamudio, and Yvona Nikitin for assistance with lab work. We thank Robin Bingham, Allison Carter, Emily Chan, Tristram Dodge, Leslie Macias, Samuel Mahanes, Julius Mojica, Lee Neighbors, Nicholas Niezgoda, Kristin Olsen, Samuel Pierce, J.T. Poole, Corinne Ried, Laurel Schumm, Jack Snow, Emily Sonenshine, Genevieve Tostevin, and William De Vries for assistance with experiment setup and data collection in the field. Jennie Reithel, Shannon Sprott, and the Crested Butte Land Trust facilitated logistics for the field experiment. We thank Kathy Ghattas, Eli Hornstein, and the Duke University greenhouse staff for their assistance planting and growing the near-isogenic lines for the field experiment; the Duke University Phytotron staff for support during the plant-soil feedback experiment; and the University of Chicago greenhouse staff for support with plant growth for chemical phenotyping. Fieldwork was conducted in compliance with United States Forest Service special use permit GUN 1120 through coordination with the Rocky Mountain Biological Laboratory.

## Data availability

The raw amplicon sequence data for the rhizosphere selection experiment and the rain exclusion field experiment are deposited in the NCBI SRA under Bioproject accession numbers PRJNA1303173 and PRJNA1276542, respectively. The phenotype data from the field experiment were previously archived on Dryad (DOI: 10.5061/dryad.18931zd4s). The glucosinolate data, phenotype data from the rhizosphere selection experiment, processed microbiome data, and all code used for analyses are available in a Zenodo repository (DOI: 10.5281/zenodo.22964897).

## Author contributions

Conceptualization: L.N.C., M.R.W. Methodology: L.N.C., M.R.W., E.B. Investigation: L.N.C., A.C., A.S., E.B., M.R.W. Visualization: L.N.C., M.R.W. Funding acquisition: L.N.C. Supervision: L.N.C., M.R.W. Writing–original draft: L.N.C., M.R.W. Writing–review and editing: L.N.C., A.S., E.B., A.C., M.R.W.

## Online Methods

### I. Characterizing variation in leaf and root glucosinolate profiles

#### Field surveys of focal populations

To assess the extent of polymorphism in glucosinolate profiles in wild populations in the study area (Figure 1b), we haphazardly selected 20-40 wild plants from each focal site. The three field sites of interest in this case are the two populations from which soil was collected for use as microbial inocula in the rhizosphere selection experiment (“RM” and “UA”) and the location where the field experiment was conducted (“SCH”). All are sites with natural wild populations of *B. stricta* near the Rocky Mountain Biological Laboratory in Gunnison County, Colorado, USA.

Initial field surveys were conducted during the 2015-2016 growing seasons, and were supplemented with additional collections with denser sampling in the three focal sites in 2024. In 2015-2016, we harvested a single leaf from each representative individual in the field for chemical phenotyping. In 2024, we collected seeds from each representative individual, and harvested a single leaf from rosette plants of the progeny grown under uniform greenhouse conditions at the University of Chicago. We stored all harvested leaves in 70% HPLC-grade methanol until processing in the lab.

#### Host genotypes in growth chamber and field experiments

The plant materials used in both experiments are near-isogenic lines (NILs) of *Boechera stricta* isolating allelic variation in a focal gene cluster, *BCMA*, which encodes enzymes controlling the biosynthesis of aliphatic glucosinolates from amino acid precursors (*33*). NILs with the *BB* genotype carry two copies of the derived allele that uses branched-chain amino acids as precursors, whereas *MM* genotypes retain the ancestral phenotype of methionine-derived glucosinolates (*63*). For simplicity, in this paper we use “glucosinolates” as shorthand for both glucosinolates *per se* and their breakdown products (e.g., isothiocyanates). There are multiple independent crossing lineages (“family lines”) nested within each NIL genotype to account for possible effects of segregating loci outside of the *BCMA* region at the time that crossing lines were established. These family lines are identified as numbers, but these numbers do not denote order or quality (e.g., line LL-15 was not generated prior to line LL-19, nor are they qualitatively different in phenotype). Full details about the crossing design used to develop the NILs are provided in past work (*30*).

Foliar glucosinolate profiles of the *BCMA* NILs are well-described, but variation in their root glucosinolate profiles was unknown prior to this study. To assess relationships between leaf and root profiles, we grew *BCMA* NILs under greenhouse conditions and collected tissue for paired leaf and root analysis. We planted seeds of *BCMA* NILs into 107-mL pots (Cone-tainers, Stuewe and Sons, Tangent, OR) containing potting soil (80% Fafard 4P [Conrad Fafard, Agawam, MA], 20% Sunshine MVP [SunGro Horticulture, Vancouver, BC]) and grew them under greenhouse conditions (18-21°C daytime temperature, 13-16°C nighttime temperature, 16h day length, watering to saturation daily, fertilizer with 300 ppm N weekly) at Duke University (Durham, NC, USA) to the juvenile rosette stage. Genotypes were spatially randomized within the greenhouse bench during plant growth. Approximately 2 months after planting, we harvested root and leaf tissue from each focal individual. We stored each tissue sample in a vial of 1.7 mL of 70% methanol. Later, the tissue samples were air-dried and weighed for use in calculation of glucosinolate concentrations.

#### Glucosinolate quantification

For both field- and greenhouse-derived tissue collections, we allowed glucosinolates to leach from tissue samples into the 70% methanol for at least 1 month. We then quantified glucosinolate profiles using established high performance liquid chromatography (HPLC) protocols (*64*). We spatially randomized all samples into 96-well plates and extracted desulfo-glucosinolates from methanol leachates. Specifically, we passed each sample through a column of hydrated DEAE-Sephadex A-25 chloride (Sigma Aldrich, St. Louis, USA) that had been equilibrated in 20 mM sodium acetate and then loaded with 50 μL of 1 mM sinigrin (Sigma Aldrich, St. Louis, USA) as an internal standard. After samples were bound to the columns, we washed the columns twice with 750 μL of 70% methanol; twice with 750 μL of diH_2_O; once with 750 μL of 20 mM sodium acetate; and twice more with 750 μL of diH_2_O. We added 30 μL of sulfatase (280U/mL; Sigma Aldrich, St. Louis, USA) to each column and incubated samples overnight. We then eluted desulfo-glucosinolates into 150 µL of HPLC-grade water.

For the 2015-2016 field collections and the growth chamber NILs, samples were separated on a Zorbax Eclipse XDB-C18 column (4.6 x 150 mm, 5 μm pore size; Agilent Technologies, Santa Clara, USA) on an Agilent 1100-series HPLC machine with a diode array detector. We separated individual desulfo-glucosinolates at 40°C on a gradient of water and acetonitrile (ACN) as follows: 6 minutes at 1.5% ACN; a 2-minute increase to 2.5% ACN; a 7-minute increase to 5% ACN; a 2-minute increase to 18% ACN; a 6-minute increase to 46% ACN; a 1-minute increase to 92% ACN; and a 5-minute decrease to 1.5% ACN (total run time: 29 minutes per sample). We identified the separated glucosinolate compounds based on their retention times and UV absorption spectra at 229 nm. The 2024 field collections were analyzed on an Agilent 1260-series HPLC with a Waters HSS T3 column (2.1 x 50mm, 1.8 µm) at 25°C. Separations used a gradient with acidified water (containing 0.1% v/v H_3_PO_4_) and acetonitrile (ACN) at 0.6 ml/min: The gradient (% ACN) was 1–5% (0–1.0 min), 5–8% (1.0–1.3 min), 8% (1.3–2.6 min), 8–92% (2.6–2.8 min) and 92% (2.8–4.75 min), followed by 3 min of re-equilibration at 1% ACN. Internal quality control tests have shown negligible differences in glucosinolate profiles using the longer- vs. shorter-running gradient and column methods (unpubl. data).

HPLC-DAD (*65*) data were imported into R using chromConverter 0.11.0 (*66*) and processed with chromatographR 0.9.0. For each dataset, chromatograms were baseline corrected by subtracting fitted baselines from each sample (detailed methods for each dataset described below). Retention time drift between runs was corrected by landmark-based parametric time warping: for each group, a polynomial warping function was fitted to landmark peaks detected from its average trace at 228 nm. Peaks were integrated by fitting a bi-exponentially modified Gaussian function with a sloped local baseline to each peak in the unwarped chromatograms at 228 nm, so that warping would not distort peak areas. Peak areas were taken from the fitted curves and group-wise warps were applied to the fitted retention times. Target compounds were assigned by retention time relative to a reference run containing every target.

The NIL dataset was baseline corrected by adaptive smoothness penalized least squares (asPLS, λ = 10^10^). In one sequence (28 runs), every compound eluted as a split peak; there, each target’s area was taken as the sum of the fitted peaks spanning both halves, which recovered the full peak area. Warping was skipped on this dataset because it did not improve the alignment. The 2015-2016 field dataset was baseline corrected using asymmetrically reweighted penalized least squares (arPLS, λ = 10^8^). Matches to 1ME GSL were rejected if their spectral correlation with the reference fell below 0.7 to avoid matches with another small non-glucosinolate peak that fell within its window. The 2024 dataset was baseline corrected using asPLS (λ = 3 × 10⁵). 2OH1ME glucosinolate was only detected in the samples from the 2024 field collections.

For all samples, we converted HPLC peak areas to relative and absolute concentrations of all detected glucosinolate compounds using established methods (*67*). Briefly, we divided raw peak areas by published relative response factors for each compound (*68*, *69*) to adjust for differential UV absorption. We then divided these corrected areas by the peak area for the internal standard sinigrin (which is not endogenously produced by *B. stricta*), and multiplied these values by 0.05 μmol, the amount of internal standard included in each sample, to calculate the absolute quantities of each compound. We divided absolute quantities by the sum of all detected glucosinolates to calculate the proportion of total glucosinolates comprised by each underlying compound. We calculated BC-ratio, the proportion of total aliphatic glucosinolates derived from branched-chain amino acid precursors, as previously described (*30*). Finally, for samples for which tissue mass was measured, we calculated absolute total quantities by the dry mass of each sample to standardize glucosinolate production by the amount of tissue included in each sample.

#### Analysis of spatial and tissue-specific variation in glucosinolate profiles

We used these processed HPLC data to answer several research questions: (Q1) What are the distributions of BC-ratio in the natural populations relevant to the growth chamber and field experiments conducted in this study? (Q2) Are leaf BC-ratio and root BC-ratio correlated in *BCMA* NILs? (Q3) How does glucosinolate profile differ between NIL leaves and roots? (Q4) Do the quantities of root-specific or root-enriched glucosinolate compounds (which have rarely been studied in *B. stricta*) differ between *BCMA* NIL genotypes?

To answer Q1, we assessed the distribution of BC-ratio expressed across the haphazardly selected wild individuals included in our field surveys in three wild populations (Figure 1B). To answer Q2, we fit root BC-ratio phenotypes in response to leaf BC-ratio phenotypes using a univariate linear model (fig. S1A). We tested the significance of this relationship using ANOVA. To answer Q3, we visualized the relative abundance of each constituent glucosinolate in each sample and identified compounds with clear patterns of differential abundance in roots vs. leaves (fig. S1B). To answer Q4, we formally modeled the concentrations of 6MTH and 4MOI3M, which are enriched in roots, as responses to the fixed effects of *BCMA* genotype, tissue type (root vs. leaf), and a random effect of plant unique ID (since root and leaf quantities were paired measurements from the same individual plants), using a linear mixed effects model (fig. S1C-D). We tested the significance of fixed effects using type III ANOVA.

### II. Rhizosphere conditioning experiment

#### Field soil collections

We sampled bulk soil from two field sites in Colorado, USA: Upper Avery (“UA”; 38.9810°, - 106.9907°; elevation ∼3,114 m) and Ross’s Meadow (“RM”; 38.9958°, -107.0137°; elevation ∼2,988 m). Prior surveys of natural populations at these two sites show that the population at Upper Avery generally produce leaf glucosinolate profiles with intermediate to high BC-ratio, while plants at Ross’s Meadow generally have low BC-ratio (Fig. 1B). We used a surface-sterilized trowel to dig up ∼1 gallon of bulk soil from each site in August 2016. Field soil was taken from the upper 15-30cm of topsoil after removing surface debris from several microsites within a ∼2m radius of each other. The focal microsites were located near the center of experimental arrays deployed earlier in 2016 (*30*) but were locations with undisturbed soil prior to sampling. We stored samples in sealed plastic bags at 4°C for use in later experiments.

#### Conditioning phase: Manipulating rhizosphere microbiomes in the greenhouse

To better control microbial communities in greenhouse experiments, we sterilized potting soil before use. Specifically, we packed Fafard 4P (Conrad Fafard, Agawam, MA) into autoclave bags and autoclaved them for 25 minutes on a dry cycle. We then removed the bags and left them to cool at room temperature, still sealed, for a minimum of 24 hours. We repeated this process twice more, autoclaving soils a total of three times with 24 hours between cycles.

We “seeded” microbial communities by inoculating sterilized potting soil with field microbes. To do this, we combined sterilized potting soil with live field soils (5% v/v) in a large, clean, surface-sterilized bin (*70*). We mixed the live soil into the potting soil thoroughly using a surface-sterilized (70% EtOH) stainless steel trowel. We prepared separate soil inocula for each of the two field soil sources. We then filled clean pots with inoculated potting soil for planting, surface-sterilizing all soil-handling tools between uses to prevent cross-contamination between starting field soil inocula.

To reduce effects of genotypic and environmental variation in initial surface microbes on seeds, we surface-sterilized them using chlorine gas. Specifically, we transferred seeds into labeled 1.5 mL Eppendorf tubes. We then placed seed tubes, with lids open, with a beaker containing 100 mL bleach inside of a 7 L desiccator chamber inside of a fume hood. We added 3 mL of HCl to the beaker of bleach and sealed the lid onto the desiccator chamber using Parafilm. We left seeds inside the sealed chamber for a minimum of 1 hour, opened the desiccator chamber (with the fume hood still closed) for 3 hours to ventilate, and then closed the lids of the tubes containing sterilized seeds.

We sowed surface-sterilized seeds into the sterilized and field-inoculated potting soil using surface-sterilized (70% EtOH) spatulas and forceps to handle seeds and re-sterilizing between pots. We grew pots in growth chambers (22°C daytime temperature; 17°C nighttime temperature; 16 h day length; 700-800 μmol light intensity; ambient humidity) for 8 weeks post-germination, watering pots to saturation daily with reverse osmosis (RO) water and fertilizing twice weekly with 0.5x Hoagland’s solution. We initially sowed 5 surface-sterilized seeds per pot, and in pots with multiple germinants, we thinned seedlings using surface-sterilized forceps (70% EtOH) to one per pot. We planted 48 pots per combination of *BCMA* NIL genotype and field soil inoculum, with the 48 replicates distributed evenly across 4 independent family lines per genotype. We also included 48 control plots per field soil inoculum treatment into which no seeds were sown. In total this yielded 192 experimental pots (2 genotypes * 2 soil treatments * 4 family lines per genotype * 12 replicates per family line) and 96 control pots (2 soil treatments * 48 replicates) for a total of 288 initial pots. Eighty-eight pots were removed from the experiment in Cycle 1 because they had no germinants to initiate rhizosphere conditioning. One additional pot (#197) was the sole pot in that genotype*soil treatment combination from its family line that had any germination; because there was no replication in this category, this pot was also removed from the experiment.

We ceased watering and fertilizing 4 days prior to the census date to make handling roots easier. We then harvested all living individuals. We used sterilized forceps to gently remove roots from the soil and shook roots gently to dislodge most bulk soil. We then placed the harvested roots, including the thin layer of rhizosphere soil in direct contact with the roots, into a 15mL Falcon tube containing sterile 1x phosphate-buffered saline (PBS). We vortexed tubes vigorously to dislodge rhizosphere soil from the roots, then removed the root tissue using surface-sterilized (70% EtOH) forceps.

At the end of the first cycle of rhizosphere selection, we used the rhizosphere-PBS slurry to immediately inoculate fresh pots containing sterilized potting soil to initiate Cycle 2. We then planted surface-sterilized seeds into the newly inoculated pots and proceeded to germinate and grow a second cycle of plants as described above. Eight weeks post-germination, we scored survival, aboveground biomass, leaf number, and height, and harvested rhizosphere soil as described above. We used the Cycle 2 rhizosphere slurries to inoculate sterilized potting soil to initiate Cycle 3 of the experiment, and germinated, grew, and measured plants as described above. At the end of Cycle 3, we harvested rhizosphere soil as described above. At the end of each Cycle, we collected aliquots of each rhizosphere for storage at -20°C until later DNA extraction and sequencing; the remainder of each rhizosphere slurry was stored at 4°C until use.

The microbial communities from each of the 199 pots were maintained independently through the end of the experiment. Of these, 74 host-conditioned rhizosphere lineages and all 96 bulk soil microbial lineages persisted through to the end of Cycle 3 of rhizosphere selection and could be included in downstream data analysis and as inocula for the reciprocal transplant experiment. The reason for additional loss of replication after Cycle 1in the host-conditioned lineages was failed germination in Cycles 2 and 3.

#### Test phase: Measuring feedback effects of rhizosphere conditioning on plant phenotype and fitness

##### Reciprocal transplant experimental design

The “test phase” of the project employed a reciprocal transplant experiment to assess the effects of host-conditioned microbiomes on plant performance. Briefly, we inoculated *BB*- and *MM-*conditioned microbiota from the end of rhizosphere selection Cycle 3 into pots containing either *BB* seeds or *MM* seeds. Although the microbiota derived from RM and UA soils slowly converged in composition during the conditioning phase, perhaps due to gradual acclimation to the autoclaved potting soil medium, the microbial communities remained distinguishable throughout the experiment and were still representative of the two natural habitats from which they were collected by the time they were used to inoculate plants in the test phase (figs. S5, S6).

To simplify the experimental design, we formed the inocula by pooling six replicate microbiota lineages within each genotype-soil source combination (*e.g., BB-*conditioned UA microbial pool). Among the 74 host-conditioned microbial lineages that persisted to the end of Cycle 3, we selected six replicate lineages within each genotype-soil source combination. When possible, we included three independent microbial lineages representing each of two family lines. If more than two family lines had three or more microbial lineages persisting to the end of Cycle 3, we used a random number generator to determine which to include in the pooled inoculum. Similarly, if a family line had more than three microbial lineages persisting to the end of Cycle 3, we used a random number generator to choose which microbial lineages to include in the pooled inoculum. In addition to the host-conditioned lineages, we randomly selected six control (unplanted or bulk soil control) lineages from each starting soil treatment to pool into inocula. In total, this created four experimental inocula (2 soil sources * 2 conditioning genotypes) and two control inocula (2 starting soil treatments * 1 non-host conditioning phase treatment [to track microbiome changes in bulk soil in absence of a conditioning plant]), each of which was pooled from six independent microbial lineages within that category. Of the 15 mL of Cycle 3 rhizosphere slurry available per sample, we used 5 mL in the pooled inoculum, yielding a total volume of 30 mL per inoculum.

After sterilizing potting soil as described above, we filled pots with soil, saturated them with 0.5x Hoagland’s solution (in RO water), and inoculated each pot with 500 μL of one of the six inocula. We inoculated 48 pots per pooled inoculum, and sowed 10 surface-sterilized seeds (as described above) into each pot. We refer to the resulting plants as “test phase” plants, and their genotypes (*BB* or *MM*, distributed evenly within each inoculation treatment) as the “test genotypes” to distinguish them from the “conditioning genotype” (also *BB* or *MM*), which is a property of the inoculum. In total, we planted 288 pots to test for feedbacks of host-conditioned microbial community composition on plant performance (6 inocula * 2 genotypes * 3 family lines per genotype * 8 replicates per family line). In addition, we included 48 control pots which contained sterilized potting soil with no inoculum added, into which we planted seeds from 8 replicates of 3 family lines of the 2 CFR-NIL genotypes (2 * 3 * 8). This yielded a total of 336 pots in the reciprocal transplant experiment.

##### Test phase plant growth and phenotype data collection

We monitored germination by counting germinants in each pot every three days for three weeks. After three weeks, ∼75% of pots had at least one germinant, and the rate of germination had plateaued. For purposes of analyzing germination success, we recorded the final number of germinants per pot after three weeks as the response variable. Then, to maximize sample size for purposes of analyzing post-germination performance and fitness, we transplanted extra germinants into pots with no seedlings, provided that the inoculum treatment and plant genotype were matched, using surface-sterilized forceps. Finally, we thinned any remaining seedlings to one per pot.

After ∼9 weeks of growth, we counted the number of leaves per plant at the end of the juvenile stage. We then vernalized plants for 8 weeks to induce reproduction (4°C daytime, 4°C nighttime, 16h day length, ambient humidity, watering to saturation with RO water as needed). After vernalization, we returned plants to normal growth conditions described above and checked pots for flowering daily for two weeks, recording the date of first flowering. After two weeks, ∼77% of individuals had flowered and the rate of new flowering had slowed; after that, we continued to census and record first flowering events at 2-3 day intervals for one additional week, at which point ∼85% of individuals had flowered. No first flowering date was recorded for 17 individuals that flowered after the final flowering time census, and for 35 individuals that died without flowering.

Under permissive greenhouse conditions, survival was close to 100%; therefore, we analyzed only the fecundity component of fitness. We censused reproductive output ∼5 weeks after the final flowering time census, by which time most buds on each plant had flowered and produced elongated, mature siliques. During the reproduction census, we measured plant height (*i.e.,* height of the longest bolting stalk); the number of total leaves produced by the end of the adult stage; the number of reproductive structures (fruits plus any late flowers that had not yet developed into fruits); and the total size of all fruits produced on each plant. To estimate total fruit size, we binned all fruits into size classes (0.5-1 cm; 1-2 cm; 2-3 cm; 3-4 cm; 4-5 cm; 5-6 cm; >6 cm) and summed the products of the number of fruits in each size bin times the lower size limit for each bin.

##### Statistical analyses

We used data from the rhizosphere reciprocal transplant experiment to answer two main research questions: (Q1) Does the effect of rhizosphere microbes on the expression of plant phenotypes and/or fitness depend on plant genotype in the conditioning generation? (Q2) What are the implications of this potential plant-soil feedback for maintenance of the *BCMA* polymorphism?

In answering these questions, we examined a suite of plant phenotype and fitness response variables: germination, juvenile leaf number, height at flowering, flowering time, adult leaf number, and the number of flowers and fruits. For Q1, we subsetted data to just the host-conditioned inocula in the test phase and modeled each plant phenotype separately in response to fixed effects of the conditioning plant genotype, test plant genotype, their interaction, and random effects of experimental block and test phase family line nested within genotype. We used linear mixed effects models for response variables that were continuous (height at flowering, flowering time, total fruit length) and generalized linear mixed effects models with a Poisson distribution for response variables that were integer counts (juvenile leaf number, adult leaf number, and number of flowers+fruits). In all models, we tested the significance of main effects using Type III ANOVA. Prior to analysis, we discovered that one family line (“L9”) had an inconsistent glucosinolate phenotype; we therefore excluded L9 plants from all analyses, for this experiment and all others described in this paper.

To answer Q2, we compared fitness across each combination of test genotype and conditioning genotype to calculate the interaction coefficient (*I_S_*), a widely-used statistic that can differentiate positive from negative feedback loops when applied to a reciprocal two-genotype experimental design (*19*):

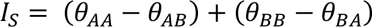

where *Θ_X_*_F_ is the estimated fitness of test genotype *X* in soil conditioned by genotype *Y.* These fitness values were estimated marginal means from the Poisson model of flower/fruit number described above. Using the emmeans package in R, we re-gridded the estimated marginal means to the original data scale (counts of reproductive structures) and then applied a custom contrast assigning weights of +1 to the self-conditioned means (*Θ_X_*_F_ where *X=Y*) and weights of -1 to the other-conditioned means (*Θ_X_*_F_ where *X≠Y).* We then calculated separate point estimates and 95% confidence intervals for *I_S_* in the two starting soils/microbial pools.

Finally, because our experimental design included non-inoculated pots and pots inoculated with non-host-conditioned (“bulk soil”) microbiomes from the test phase, we also used these data to ask two further questions: (Q3) Do plant phenotypes and/or fitness differ between inoculated vs. non-inoculated soils? (Q4) Do plant phenotypes and/or fitness differ for test plants grown in soil inoculated with host-conditioned rhizosphere microbiomes vs. bulk soil microbiomes? These questions assess general effects of the soil microbiome on the test plants, agnostic to genotype and starting soil pool in the conditioning generation. We answered these questions using the same model structures described above for Q1, assessing variation in germination, juvenile leaf number, height at flowering, flowering time, adult leaf number, and the number of flowers and fruits in response to the fixed effects of inoculation treatment (Q3: any live inoculum vs. none; Q4: live inoculum from a host-conditioned rhizosphere vs. live inoculum from bulk soil), test plant genotype, and their interaction, and random effects of experimental block and family line.

Analyses for Q1 and Q2 are the main hypotheses tested in this paper and are reported in the main text. Analyses for Q3 (table S8a) revealed that the probability of germination was marginally higher in pots inoculated with microbes than pots containing only sterilized potting soil (fig. S7A) and that the expression of genetic variation for adult leaf number was strongly responsive to inoculation, with leaf number suppressed by live inoculum for the *BB* genotype and slightly increased for *MM* (fig. S7B). No other traits or fitness components differed depending on the live vs. sterile potting conditions. Analyses for Q4 showed that the total number of fruits and flowers was higher for both genotypes when they were grown in inocula from bulk soil than when grown in inocula with host-conditioned rhizosphere microbes (table S8B; fig. S7C).

#### Characterization of conditioning phase microbial lineages

##### Amplicon sequencing of bacterial and fungal barcode genes

We used bacterial and fungal barcode gene sequencing to investigate how host plant genotype and phenotype influenced microbial community composition during the rhizosphere selection experiment. Samples derived from the greenhouse rhizosphere selection experiment (Conditioning phase) were collected as slurries (in 1x PBS) separated from roots while preparing inocula for the Test phase reciprocal transplant experiment. Slurry subsamples were centrifuged at 4,350x g for 11 minutes, the supernatant was discarded, and pellets were resuspended in 900 μL lysis buffer (1M Tris pH=8.0, 0.5 mM NaCl; 10 mM EDTA) then transferred to randomized locations in 96-well plates containing 450 μL of garnet particles (∼2 mm diameter, Biospec Products, Bartlesville, OK, USA) per well. The same randomization scheme was maintained throughout DNA extraction and library preparation. Each 96-well plate also contained a negative control with no host plant tissue and a positive control containing a mock microbial community containing known microbial strains in known relative abundances (Zymo Research, Irvine, CA).

At the University of Kansas, we generated libraries to sequence bacterial and fungal amplicons following an established two-PCR-step protocol (*71*). Briefly, we used the 515f-806r primer pair (*72*, *73*) in the first PCR to amplify the V4 region of the bacterial/archeal 16S rRNA gene, and the ITS1f-ITS2 primer pair (*74*, *75*) to amplify the fungal ITS1 region. We amplified the 16S-v4 and ITS1 regions in separate reactions. When amplifying 16S-v4 from root (but not rhizosphere) samples, we added 0.15μL of 100μM peptide nucleic acid per 10 μL reaction volume to block amplification of host plastid sequences (*76*). We added dual-indexed i7/i5 Illumina adapters in the second PCR.

We pooled amplicon libraries and sent them to the Duke University core sequencing facility in summer 2020. Pooled libraries were size-selected to remove fragments smaller than 300 bp using Pippin Prep, and then sequenced (250 bp, paired-end) using two lanes of a NovaSeq SP flow cell. In May 2024, we re-made libraries for select samples that had an inadequate amount of data using the same protocols described above, removed fragments <300 bp using Pippin Prep, and then re-sequenced them on the NextSeq2000 platform with P1 reagents (250 bp, paired-end) at the University of Kansas Genome Sequencing Core.

##### Sequence data processing

Our established bioinformatic pipeline (*71*) was used to process raw FASTQ files from all sequencing runs described in this paper, with minor modifications depending on the depth and quality of reads. First, we used trimmomatic (*77*) to remove NextSeq adapter contamination, allowing 2 seed mismatches and using palindrome and simple clip thresholds of 30 and 10, respectively. The minimum adapter length for palindrome mode was 2 and sequences <100 bp in length were discarded. Second, we simultaneously sorted 16S reads from ITS reads and removed gene primers using cutadapt (*78*), requiring an overlap of 5 bases, an error rate of 0.15 or lower, and zero ambiguous bases. The --nextseq-trim=20 option was used during this step. Third, we concatenated the first 500 reads from each sample into a single FASTQ file which we used to estimate the error rate with the learnErrors() function in DADA2 (*79*). This was done separately for each barcoding gene (16S-v4 and ITS1) and each sequencing run. Fourth, we used DADA2 to perform additional quality control, dereplication, and denoising, discarding forward and reverse reads with more than 2 or 3 expected errors, respectively. For 16S only, forward and reverse reads were truncated at 220 bp and 200 bp, respectively. We used the core DADA2 functions and the previously estimated error rates to dereplicate and denoise all reads, merge them into a single amplicon sequence variant (ASV) table, and remove chimeras using the consensus method. This was done separately for each sample to allow parallel processing. For samples that were re-sequenced, the data from the two sequencing runs were processed separately using the same pipeline (including software versions and parameters) through ASV table generation, at which point the ASV tables were merged. However, error rates used for denoising were estimated separately for the two sequencing runs. Finally, the ASV tables from all samples were merged to yield one bacterial and one fungal table, and taxonomy assignments were made using the RDP Classifier trained on the RDP database (v19) for bacteria (*80*) or the UNITE database for fungi (*81*).

The resulting ASV and taxonomy tables were loaded into R for processing using the phyloseq package (*82*). ASVs with ambiguous taxonomy at the Kingdom level and those classified as host sequences were removed. We used the microfiltr software (*83*) to perform additional filtering based on ASV relative abundances both within and across samples, and prevalence across the dataset, with the filtering parameters tailored to each dataset to maximize both data quality and quantity. We implemented a minimum within-sample relative abundance cutoff of 0.075%, which removed 93% of contaminant ASVs from our positive controls while retaining >80% of all data and all of the expected ASVs. We removed samples with fewer than 1,000 usable (non-host, non-contaminant) reads. Alpha diversity was estimated for each sample as the Shannon index using the vegan package (*84*). Next, to maximize data retention while removing ultra-rare and non-reproducible sequences, we removed bacterial ASVs that represented <0.005% of the total dataset by relative abundance and were observed in <3% of samples. The analogous cutoffs for fungi were 0.5% within-sample relative abundance, 0.01% across-sample relative abundance, and 0.25% prevalence. After quality control, the final dataset included 1,056 bacterial ASVs and 445 fungal ASVs. The mean numbers of observations per sample were 113,720 (s.d. = 157,096) for bacteria and 20,207 (s.d. = 43,940) for fungi.

##### Statistical analysis of microbiome data from conditioning experiment

Due primarily to uneven germination throughout the conditioning phase, the final sample sizes were *N=*70 for *BB-*conditioned rhizospheres, *N*=217 for *MM-*conditioned rhizospheres, and *N*=272 bulk soil (unconditioned) controls. To minimize the impact of the unbalance between *BB* and *MM* rhizospheres, we partitioned variance using ANOVA with Type III sums of squares where possible, noting that this approach yields conservative results for hypothesis tests of marginal main effects (*85*). We applied centered log-ratio (CLR) transformations to raw ASV counts to both resolve compositionality and normalize ASV abundances (*86*, *87*). Transformations were applied using the aldex.clr() function from the ALDEx2 package (*88*); the median values from 128 Monte Carlo simulations per sample were used as the inputs for downstream analyses. The Euclidean distance matrix based on the resulting CLR-transformed ASV tables (also known as the Aitchison distance matrix) was the input for ordination-based analyses. Because the process of transferring soil microbiota between pots had an outsized impact on community composition that was unrelated to accumulating effects of host influence (fig. S5) we considered passaging cycle to be a nuisance variable. We statistically controlled for the effects of both cycle and log-transformed sequencing depth by partialling them out during ordination-based analyses and including them as nuisance covariates in univariate models of microbiome features.

To test the hypothesis that *BB-*conditioned and *MM-*conditioned rhizosphere communities differ in alpha diversity, we fit a linear mixed-effects model of Shannon diversity (*e*^Shannon_metric^) with the fixed-effect predictors Soil (UA or RM), *BCMA* (*BB* or *MM*), Cycle, and all possible interactions between them; log-transformed sequencing depth as a continuous nuisance covariate; and experimental block and family–a genetic lineage nested within *BCMA* genotypes–as random intercept terms. However, one or both random effects were removed from the model if they caused a singular fit. Models were fitted using the lmer() function in the lme4 package (*89*) and formal hypothesis testing was conducted using ANOVA with Type III sums of squares in the lmerTest (*90*) or car (*91*) package depending on whether a mixed-effects or fixed-effects model was used. Likelihood ratio tests of reduced models were used to assess statistical support for random effects. Marginal effects of predictor variables were estimated using the emmeans package (*92*).

To test the hypothesis that *BB-*conditioned and *MM-*conditioned rhizosphere communities differ in taxonomic composition, we used the capscale() function of the vegan package to conduct a partial constrained ordination of Aitchison distance matrices (*93*, *94*). The nuisance variables Cycle and log-transformed sequencing depth were partialled out; the constraining variables were Soil, *BCMA*, and their interaction. We used redundancy analysis (*95*) to assess statistical support for each of the constraining variables.

To assess the sensitivity of individual ASVs to host *BCMA* genotype, we modeled the CLR-transformed counts of each as described above for Shannon diversity. We included both experimental block and plant family as random intercept terms in the default model, but dropped one or both if needed to prevent singular model fits. In cases where both were dropped, we fit fixed-effects linear models and conducted the Type III ANOVA using the car package. P-values were adjusted for multiple comparisons using the Benjamini-Hochberg false discovery rate.

### III. Field experiment

#### Experimental design and data collection

We utilized an existing field experiment testing for effects of drought and herbivore pressure on selection on *BCMA* to determine whether NIL host genotype effects on the microbiome were also detectable in field environments. Full details on the methods of the field experiment were published previously (*31*). Briefly, in 2015, replicate individuals spanning multiple family lines of each *BCMA* NIL genotype were germinated in non-sterilized potting soil and reared through the juvenile rosette stage in the Duke University greenhouse. After ∼10 weeks of juvenile growth, rosettes were shipped to the Rocky Mountain Biological Laboratory in Colorado and transplanted into a common garden in Schofield Townsite, in a subalpine meadow reflecting typical *B. stricta* habitat. For the first year following transplanting (2016), experimental plants experienced ambient environmental conditions in the common garden. For the subsequent three years (2017–2019), surviving individuals were assigned into experimental treatments where drought stress was manipulated using rainout shelters (applied at the block level). We collected data on survival, growth, reproduction, and herbivore resistance from 2016-2019, which encompassed the full life span for ∼85% of transplanted individuals (*31*). The full field experiment was planted in two temporal cohorts, one planted in fall 2015 and the other planted in fall 2016; in this study, however, we only used samples from the 2015 planting cohort to control for effects of plant age on the root microbiome (*45*).

Fecundity data from this experiment is already published and archived (*31*, *96*). We leveraged this dataset to calculate total lifetime fecundity for the plants from which we sampled root and rhizosphere microbiomes in 2019 (methods below). We measured total lifetime fecundity as the sum of total fecundity (number of fruits multiplied by average fruit length) in each year after planting from 2016-2019. In *B. stricta*, seed set is strongly and linearly correlated with fruit length. In this experiment, the majority of lifetime reproductive fitness was achieved in years 1 and 2 after transplanting, although a small number of plants continued to produce fruits and seeds into years 3 and 4.

#### Field root/rhizosphere collection and microbiome characterization

In August 2019, we uprooted all surviving plants from the cohort planted in 2015 using a hand spade and removed loosely-adhered soil by gently flicking with gloved hands, which were rinsed clean between plants, until no more soil was falling from the roots. Entire root systems with adhering rhizosphere soil were stored in 50 mL conical tubes on dry ice until they could be transferred to a -20°C freezer. Once all root systems were collected, we transported them on dry ice to the University of Kansas, where we separated rhizosphere soil from roots by vortexing tissue in 900 μL of lysis buffer (1M Tris pH=8.0, 0.5 mM NaCl; 10 mM EDTA) for 15 seconds and then transferring the rhizosphere suspension by pipet into a 2 mL deep-well plate filled with 2.0 mm garnet particles (0.45 mL garnet per well). After the rhizosphere soil was removed, we transferred the associated root samples into cluster tubes with 2 stainless-steel ball bearings (5/32” diameter), froze them at -80°C overnight, and then freeze-dried them for 48 hours. The lyophilized roots were then powdered using a HT Lysing Homogenizer (OHAUS, Parsippany, NJ, USA). Finally, we resuspended the root powder in 800 μL of lysis buffer and transferred the resulting suspension by pipet into deep-well plates filled with garnet particles. From that point, the rhizosphere and root samples in garnet plates were processed as described above (see Amplicon sequencing of bacterial and fungal barcode genes) except that for bacterial library preparations from root samples, we added PNA PCR blockers (*76*) to reduce amplification of host plastid genes. Both bacterial and fungal libraries were generated at the University of Kansas and then sequenced on the Illumina NovaSeq (250 bp paired-end) platform at the Duke University core sequencing facility. Raw FASTQ files were processed using the bioinformatic pipeline described above (see Sequence data processing).

We filtered the resulting ASV tables using a minimum cutoff of 0.03% within-sample relative abundance, which removed all contaminant ASVs from our positive controls while retaining all of the expected ASVs. Next, we removed bacterial ASVs that represented <0.005% of the total dataset by relative abundance or were observed in <1% of samples. The analogous cutoffs for fungi were 0.05% within-sample relative abundance, 0.025% across-sample relative abundance, and 2% prevalence, which maximized data retention while removing non-reproducible sequences. After quality control, the final dataset included 1,719 bacterial ASVs and 360 fungal ASVs. Overall, 81.5% of the bacterial observations and 66.2% of the fungal observations were retained after filtering. The mean numbers of observations per sample were 121,934 (s.d. = 134,805) for bacteria and 45,744 (s.d. = 25,658) for fungi. Finally, we used ALDeX2 (*88*) to apply a centered log-ratio (CLR) transformation to the raw ASV counts; the median value of 128 Monte Carlo instances was used as the final CLR value for each ASV in each sample.

#### Statistical analysis of microbiome data from field experiment

The final quality-filtered bacterial dataset included *N*=180 samples, comprising paired rhizosphere and root endosphere (hereafter described as “root”) samples from 40 *BB* plants and 50 *MM* plants. Of these, 19 *BB* plants and 25 *MM* plants were under rainout shelters, while the remainder were in the control group. One fungal sample with fewer than 1,000 usable (non-host, non-contaminant) reads after quality control was removed from the dataset, for N=179.

##### Identifying the sources of microbiome variation in the field

First, we modeled alpha diversity (*e*^Shannon^ ^index^) using *BCMA* genotype, compartment, rainout treatment, and all interactions between them as fixed predictors with genetic family (nested within *BCMA* genotype) and plant unique ID as random intercept terms; log-transformed sequencing depth was also included as a covariate. We excluded experimental block from the model because it explained no variance and caused a singular fit. Residuals were inspected visually and variance inflation factors were checked using the car package (*91*) to confirm that assumptions of homoscedasticity and noncollinearity were met. We tested statistical significance using Type III ANOVA using default parameters in the lmerTest package (*90*). Estimated marginal means were calculated using the emmeans package (*92*).

Second, to examine whether *BCMA* effects on ASV relative abundances were equally strong in the root endosphere and the rhizosphere, we fit the same linear mixed-effects model to the CLR-transformed count of each ASV and used the contrast() function of emmeans to extract the estimated marginal mean log fold change between genotypes for each compartment. We used Spearman correlation to assess the relationship between ASVs’ responses to *BCMA* in the root endosphere and the rhizosphere.

Third, to identify experimental variables influencing overall community composition, we used Canonical Analysis of Principal Coordinates (CAP) (*93*) based on the matrix of Aitchison distances between samples (*86*). We used the vegan package (method ‘capscale’ with Euclidean distance) to ordinate the CLR-transformed ASV table. Constraining variables were *BCMA* genotype, compartment, rainout treatment, and all interactions between them; the log-transformed sequencing depth was partialled out to remove its influence prior to ordination. We assessed the statistical significance of the constraining variables using the anova.cca() method from vegan (*84*). To aid with biological interpretation of constrained axes, we visualized the effect of each constraining variable by fitting sample scores on each constrained axis to the same univariate linear mixed-effects model that we used for alpha diversity (described above) and plotting the resulting estimated marginal means.

##### Estimating natural selection on heritable microbiome variation

Because we could only sample microbiomes from plants that survived to the end of the four-year-long experiment, we analyzed only the fecundity component of evolutionary fitness. To investigate whether heritable (*BCMA*-sensitive) microbiome variation was associated with host fitness, we conducted a phenotypic selection analysis (*40*, *97*). Past work had established that *BCMA* genotype influences *B. stricta* fitness in a way that depends on water availability (*30*); here, our goal was to determine whether root and/or rhizosphere microbiota mediated the effect of *BCMA* on lifetime fecundity in the field. To do this, we used structural equation modeling (SEM) to simultaneously analyze the direct effects of *BCMA* on fecundity and the indirect effects *via* its influence on each of the seven constrained ordination axes (CAP1-7). To isolate heritable microbiome variation and to allow for microbial fitness effects to differ between compartments and between treatments, we analyzed each compartment-treatment combination separately. First, we calculated the lifetime female fecundity of each plant as: 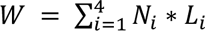 where *N_i_* is the number of fruits produced in year *i* and *L_i_* is the length in mm of the average fruit in year *i.* This value is an established proxy for seed output or fecundity in *Boechera stricta* (*30, 31, 67*). To calculate the relative lifetime fecundity of each plant (*w*), we divided the absolute fecundity (*W*) by the mean *W* of all plants in the same treatment (*40*).

The resulting *w* was used as the response variable for the SEM, fit using the lavaan package version 0.6-18 (*98*). The model included paths from *BCMA* to each of the mediating variables (CAP1-7), from *BCMA* directly to *w*, and from each of CAP1-7 to *w. BCMA* was re-coded as a dummy variable with levels (*MM*=1, *BB*=0). Residual pairwise covariances among CAP1-7 were allowed, making the path coefficients equivalent to those obtained through multiple regression (*48*). Thus, the path connecting each mediator to *w* is the directional selection gradient (*β*) on that CAP axis controlling for all others and for the direct effect of *BCMA* (*40*) expressed as units of relative fitness per unit of CAP score. Finally, we calculated the indirect effect of *BCMA* on fitness mediated through a given CAP axis as the product of the two corresponding path coefficients, in units of relative fitness. Both per-axis and summed indirect effects were computed, along with the direct effect of *BCMA* on *w*. The model was just-identified (df = 0), rendering global fit indices uninformative; however, this does not affect inference on individual path coefficients or indirect effects, for which we computed bias-corrected and accelerated confidence intervals (*99*) based on 1,000 bootstrap replicates, of which >99% converged.

##### Estimating directional selection on bacterial relative abundances in the root

Next, we used the principal component regression back-projection technique of Chong *et al.* (*39*), which allows the selection gradients estimated on ordination axes to be projected back into the original variable space (in this case, CLR-transformed ASV relative abundances). This method is preferable to direct estimation of selection on individual taxa because the use of orthogonal predictors reduces multicollinearity and thus stabilizes the selection gradient estimation. Because we used constrained axes from a Canonical Analysis of Principal Coordinates (*93*) on subsetted data, our predictors were not strictly orthogonal. Our decision to use CAP rather than traditional PCA was biologically motivated, because selection gradients are only evolutionarily meaningful for heritable traits and restricting our analysis to axes of microbiome variation that distinguish host genotypes ensured that the back-projected selection gradients met this criterion. Although the CAP axes were constrained on BCMA*Compartment*Treatment interactions, selection gradients were calculated within each compartment-treatment subset, leaving only the variation attributable to BCMA. We reasoned that this crucial biological interpretability justified the tradeoff of imperfect orthogonality. Additionally, we found that the degree of collinearity among the CAP axes was modest relative to what would be expected among the thousands of underlying taxa that can show substantial ecological covariance structure in plant-associated microbiomes (*100*), suggesting that this approach still provided substantial improvement over direct estimation of selection gradients on microbial relative abundances.

From the SEM described above, we obtained directional selection gradient estimates for each CAP axis (*β*, the path coefficient linking each axis to relative fecundity) in the root endosphere compartment separately for the control and drought treatments. This choice was biologically motivated: microbial effects on host phenotype often depend on environmental contexts such as abiotic stressors (*35*) and could reasonably be expected to vary between compartments as well. For each compartment-treatment combination, the resulting vector of regression coefficients A was back-projected into ASV space as β = E × A, where E is the matrix of ASV loadings on the heritable CAP axes (*39*). Uncertainty in per-ASV selection gradients was propagated from the model variance-covariance matrix as Var(β) = E × Var(A) × Eᵀ, and 95% confidence intervals were constructed as β ± 1.96 × SE(β). For analyses of categorical selection patterns (positive, negative, or zero), ASVs were considered to have a non-zero selection gradient if their 95% CI excluded zero.

Finally, we investigated patterns in selection on the relative abundances of individual bacterial ASVs in the root endospheres of droughted plants. First, we partitioned variance in ASV selection gradients among higher-level taxa by fitting the random-effects model:

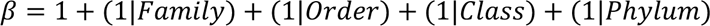

using restricted maximum likelihood (REML), weighted by the inverse of the standard error for *β*. Higher-level taxa with fewer than 5 representative ASVs were excluded from this analysis. Because the taxonomic levels are strictly hierarchical and therefore implicitly nested, we conducted sequential likelihood ratio tests by removing one term at a time, from most specific to broadest, and comparing the fits of pairs of reduced *vs.* full models. For example, statistical support for organization of selection gradients by Class was obtained by comparing the models *β* = 1 + (1|*Class*) + (1|*P*ℎ*ylum*) and *β* = 1 + (1|*P*ℎ*ylum*). These analyses were conducted using the lme4 package. We then estimated the mean directional selection gradient on ASVs within each taxon by fitting the linear models *β* ∼ *Family*, *β* ∼ *Order*, *β* ∼ *Class*, and *β* ∼ *P*ℎ*ylum*, each weighted by the inverse of the standard error for *β*. The estimated marginal means and 95% confidence intervals from these models were plotted for comparison among taxa.

We also tested whether selection gradients on ASV relative abundances in the root endospheres of droughted plants were correlated with (1) log fold changes in abundance due to drought treatment, (2) log fold changes in abundance between *BB* and *MM* plants, and (3) selection gradients on relative abundances in the root endospheres of control plants. The log fold changes were contrasts of estimated marginal means from the linear mixed models of CLR-transformed ASV abundances described previously. To propagate uncertainty in these estimates into the correlation analysis, we generated 10,000 Monte Carlo simulated datasets in which the value of each variable for each ASV was drawn independently from a normal distribution with mean equal to its point estimate and standard deviation equal to its standard error. We calculated Spearman’s rank correlation coefficient (ρ) for each simulated dataset and report the mean ρ across simulations, with the 2.5th and 97.5th percentiles as a 95% confidence interval.

---

**Figure S1.**
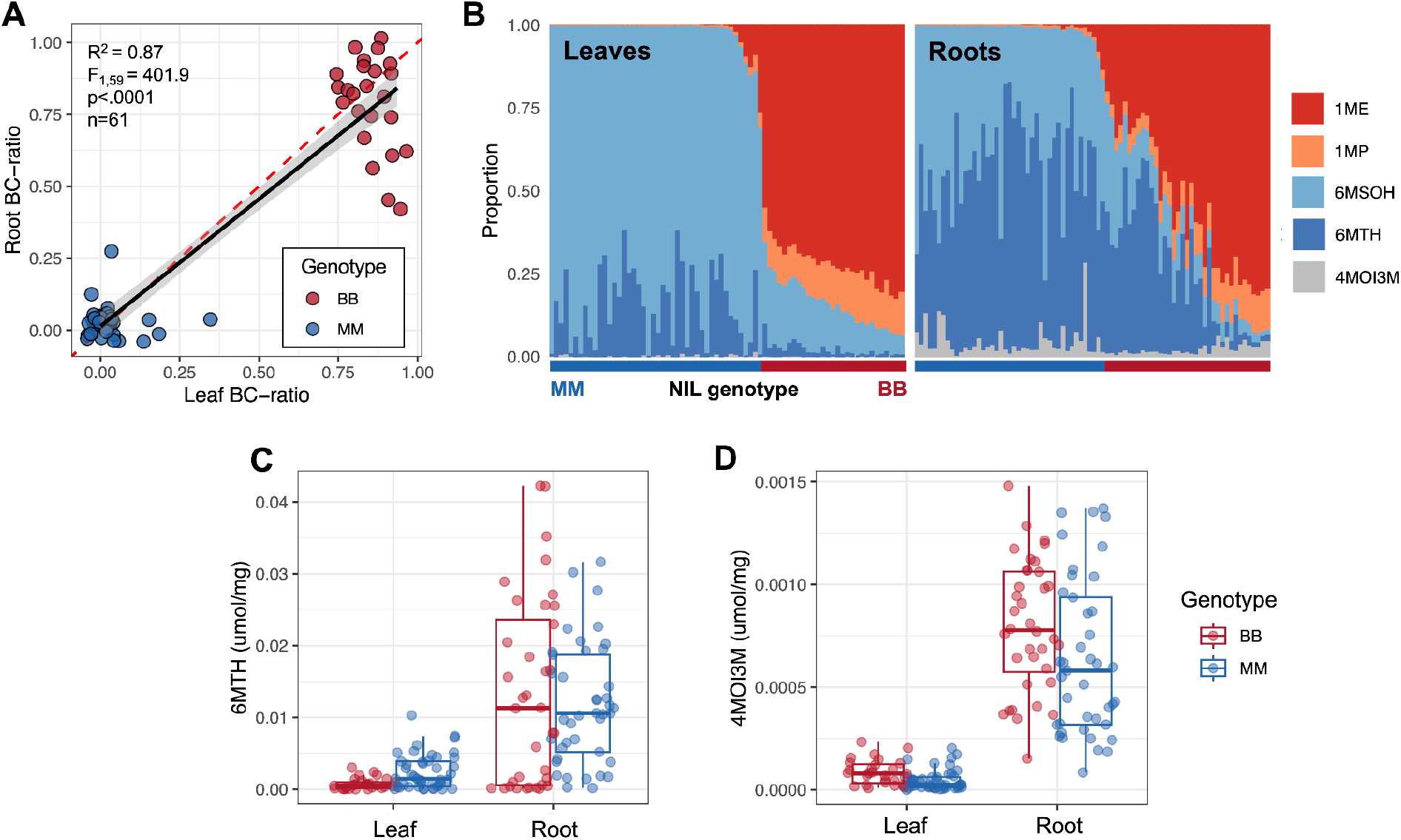
Relationships between root and leaf glucosinolate profiles in the *BCMA* NILs. (**A**) BC-ratio is strongly correlated across leaves and roots in near-isogenic lines homozygous for alternative alleles at *BCMA* (r=0.87, *F*_1,59_ = 401.9, *P*<0.0001). (**B**) *BCMA* genotype has a major effect on BC-ratio (the proportion of aliphatic glucosinolates derived from branched-chain amino acids) and overall glucosinolate composition in both leaves and roots. 6MTH (a Met-derived aliphatic glucosinolate; dark blue bars) and 4MOI3M (a Trp-derived indolic glucosinolate; gray bars) are more common in roots than leaves. Stacked colors in each bar represent the proportion of total glucosinolates comprised by each underlying compound present in the overall profile, with warm colors showing branched-chain aliphatic GLS, blues showing unbranched (Met-derived) GLS, and gray showing the indolic compound. Each bar represents an individual plant (with multiple biological replicates per genotype and family line). Samples are grouped by tissue type. The horizontal, colored bar at the bottom links each vertical bar to its genotype (blue: *MM* [homozygous for Met-GLS]; red: *BB* [homozygous for BC-GLS]). Bars are ordered left to right from lowest to highest BC-ratio. Also note that the branched-chain compound 2OH1ME, which is sometimes present in *B. stricta* leaves, was absent in this panel of samples. **(C-D)** The absolute amounts of (**C**) 6MTH and (**D**) 4MOI3M produced in roots are significantly higher than the amounts produced in leaves for both *BCMA* NIL genotypes (6MTH: main effect of tissue type, *F*_1,145_ = 39.21, *P* <0.0001; 4MOI3M: main effect of tissue type, *F*_1,145_ = 122.42, *P* <0.0001; *P*>0.05 for effects of genotype and genotype*tissue interaction).

**Figure S2.**
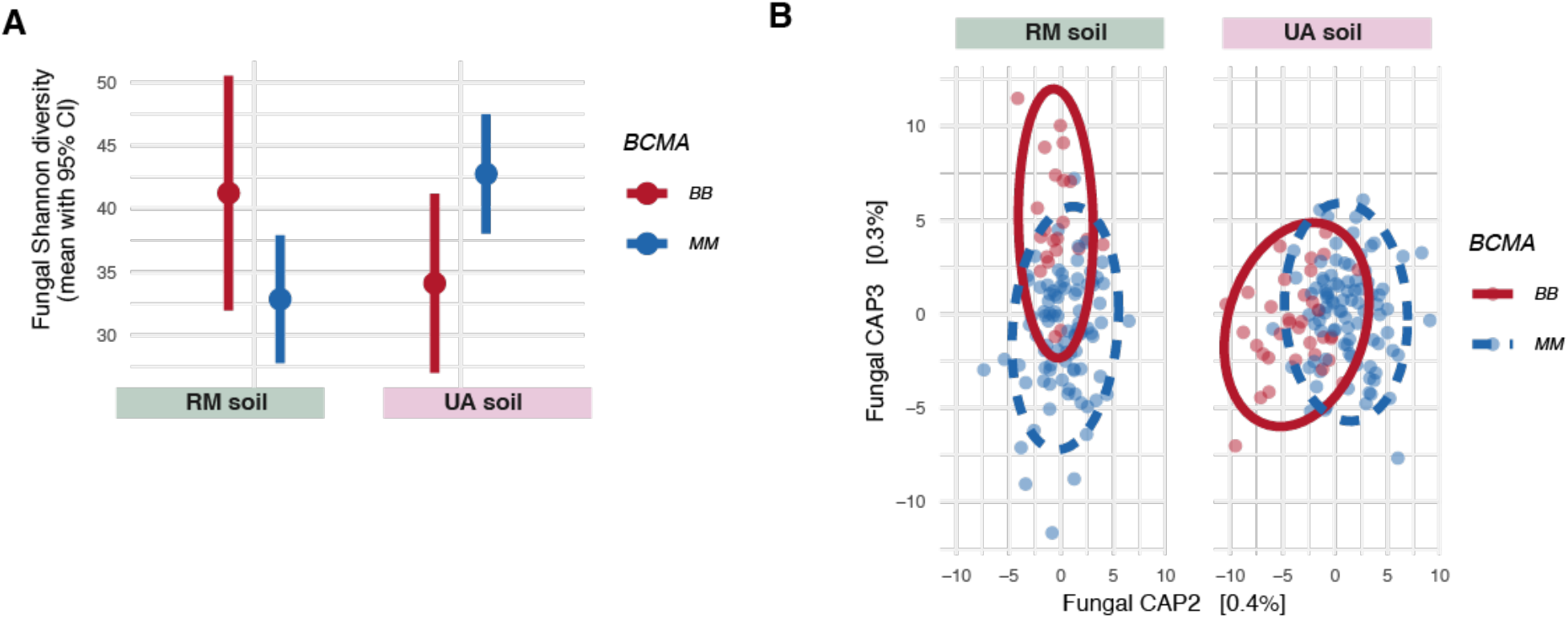
In the growth chamber, fungal rhizosphere communities conditioned by near-isogenic lines with contrasting *BCMA* alleles differed in alpha diversity but not taxonomic composition. Results are averaged across three cycles of conditioning. (A) Estimated marginal mean alpha diversity (*e*^Shannon^ ^index^) of rhizosphere fungal communities conditioned from two starting soils. Statistical support is provided in table S1. (**B**) Constrained ordination of Aitchison distances among fungal rhizosphere communities conditioned from two starting soils, with 95% confidence ellipses. Statistical support from redundancy analysis is provided in table S1.

**Figure S3.**
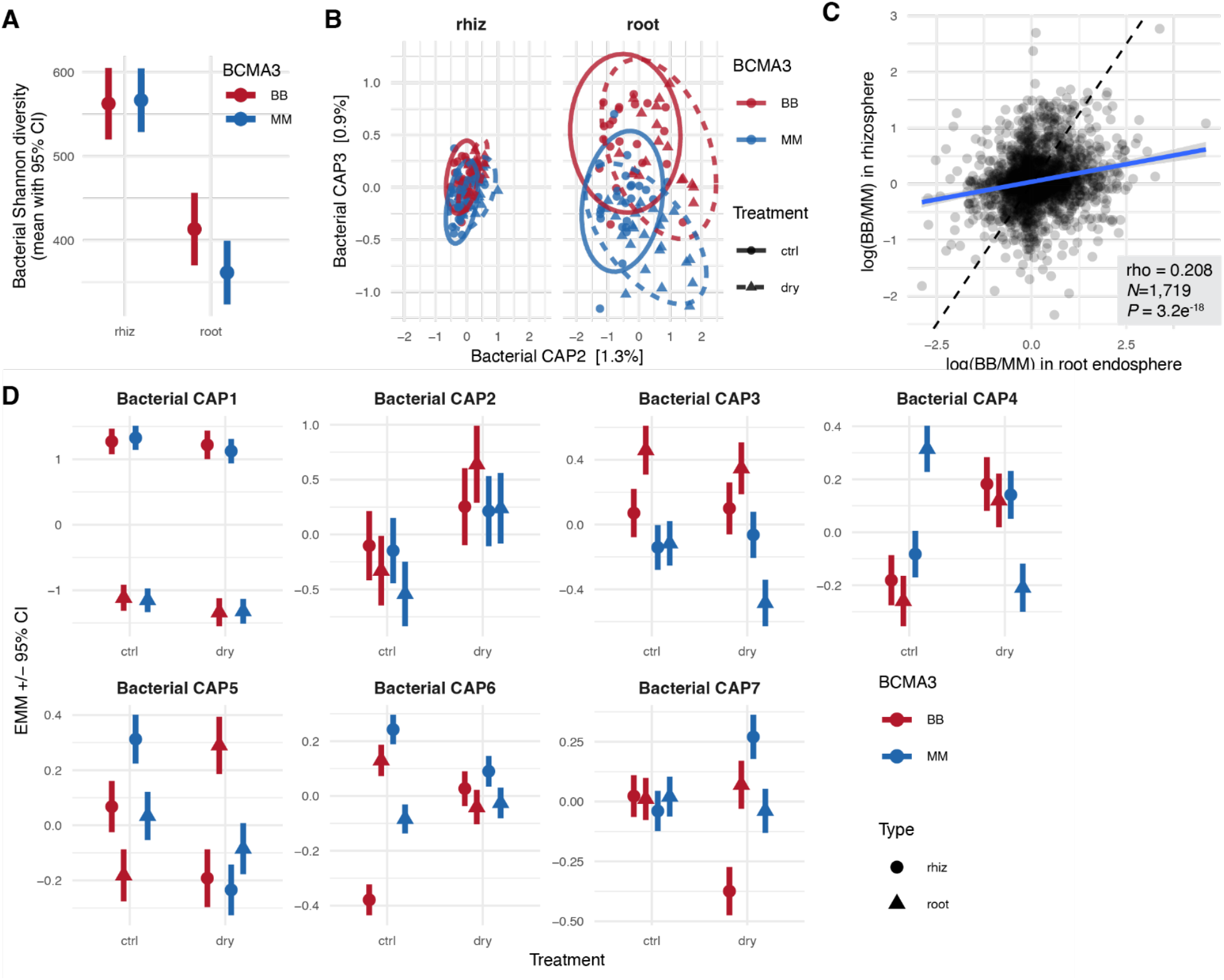
Field-grown *BCMA* near-isogenic lines assemble distinct bacterial communities in both the rhizosphere and the root endosphere. **(A)** Estimated marginal mean alpha diversity (*e*^Shannon^ ^index^) of bacterial rhizosphere communities. **(B)** Constrained partial ordination of Aitchison distances among bacterial rhizosphere communities, with 95% confidence ellipses. The effect of sequencing depth was partialled out, and the constraining variables were *BCMA*, sample type (rhizosphere *vs.* root), Treatment (drought *vs.* control), and their interactions. **(C)** The estimated marginal mean log fold change in abundance between *BB* and *MM* genotypes is plotted for each bacterial ASV in the root endosphere (horizontal axis) and the rhizosphere (vertical axis). The line of best fit is shown in blue; the dashed line represents y=x, the expectation if *BCMA* effects were equally strong in the two compartments. **(D)** Estimated marginal means of each constrained axis of bacterial taxonomic composition for each genotype, sample type, and precipitation treatment. Statistical support from linear mixed models (for alpha diversity) and term-wise redundancy analysis (for community composition) is provided in table S4. The analogous plot for fungi is shown in fig. S4.

**Figure S4.**
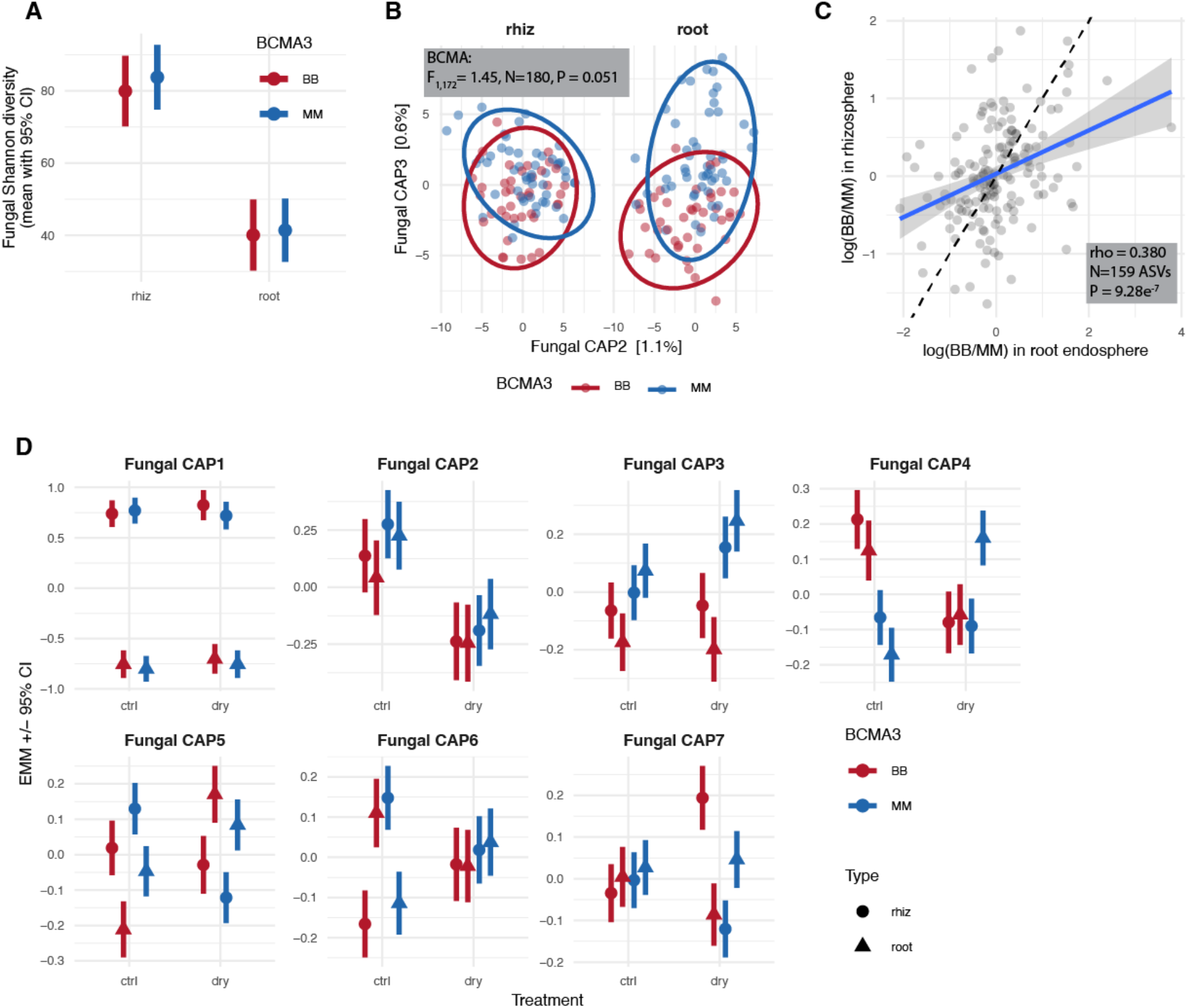
Field-grown near-isogenic lines segregating for the *BB* and *MM* alleles at the *BCMA* locus assemble distinct fungal communities in both the rhizosphere and the root endosphere. **(A)** Estimated marginal mean alpha diversity (*e*^Shannon^ ^index^) of fungal rhizosphere communities was the same between *BCMA* NILs. **(B)** Constrained partial ordination of Aitchison distances among fungal rhizosphere communities, with 95% confidence ellipses. The effect of sequencing depth was partialled out, and the constraining variables were *BCMA*, sample type (rhizosphere *vs.* root), Treatment (drought *vs.* control), and their interactions. **(C)** The estimated marginal mean log fold change in abundance between *BB* and *MM* genotypes is plotted for each fungal ASV in the root endosphere (horizontal axis) and the rhizosphere (vertical axis). The line of best fit is shown in blue; the dashed line represents y=x, the expectation if *BCMA* effects were equally strong in the two compartments. **(D)** Estimated marginal means of each constrained axis of fungal taxonomic composition for each genotype, sample type, and precipitation treatment. Statistical support from linear mixed models (for alpha diversity) and term-wise redundancy analysis (for community composition) is provided in table S4. The analogous plot for bacteria is shown in fig. S3.

**Figure S5.**
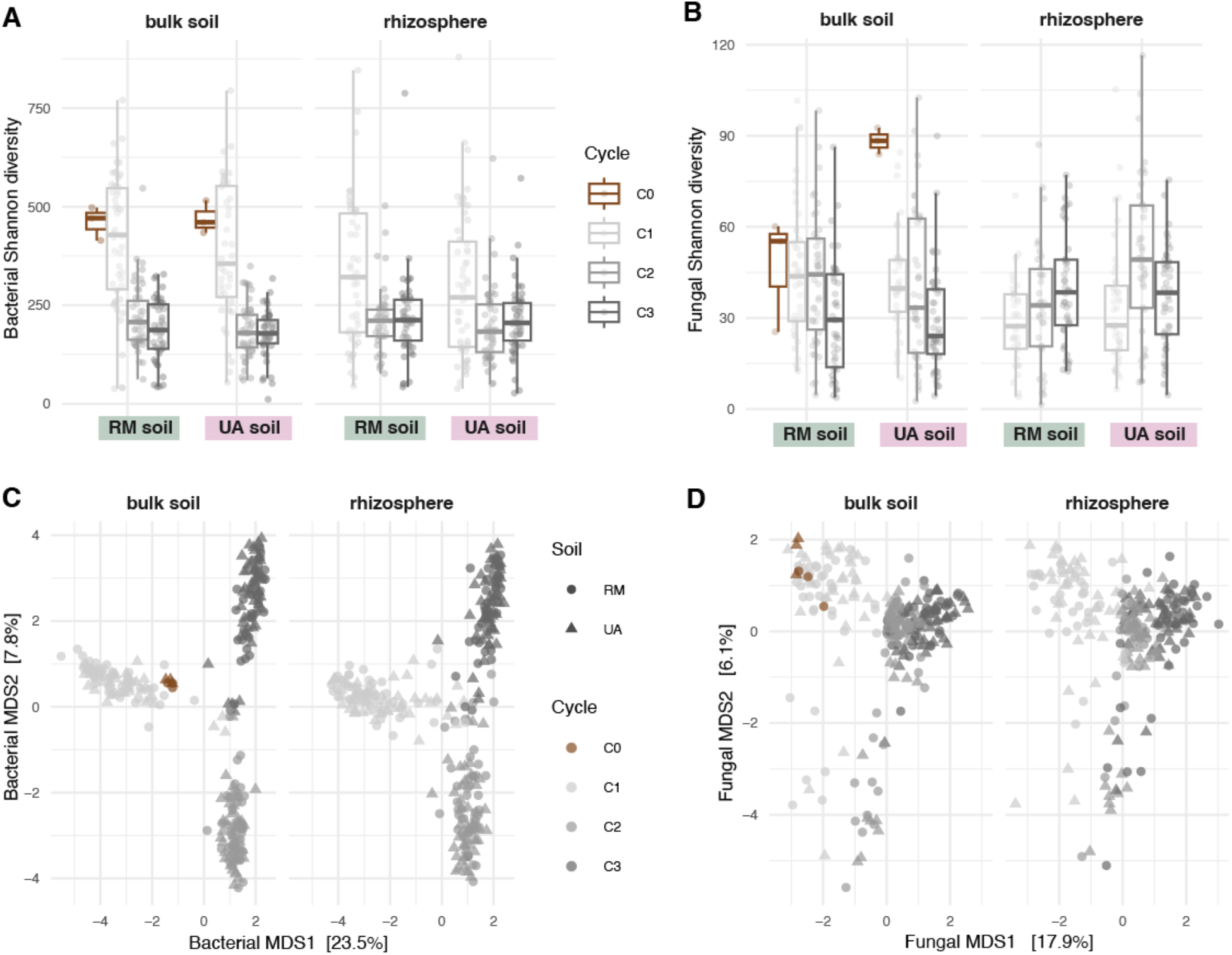
Changes in alpha-diversity (A-B) and community composition (C-D) beginning with field-collected soils (“C0”) through three cycles of passaging into sterile potting soil background (“C1-C3”). Bulk soil samples in C1-C3 were from unplanted pots; rhizosphere samples were pots planted with *Boechera stricta*. The process of passaging natural soil microbiota into the autoclaved potting soil background altered the diversity and composition of both the bacterial and fungal communities relative to the original field soils. For bacteria, a major shift in community composition separated cycle 1 from cycles 2 and 3 (A,C), suggesting that microbiome transfer via slurry (initiated in cycle 2) retained bacteria less effectively than transfer via solid soil (cycle 1). In contrast, fungal communities changed linearly over the cycles (B,D). Community dispersion declined throughout the experiment for both bacteria and fungi, indicating a homogenizing effect of the experimental conditions. These cycle effects were dominant in both the rhizosphere (planted pots) and bulk soil (unplanted pots), indicating that they primarily reflect artifacts of the passaging method rather than an accumulating effect of host presence.

**Figure S6.**
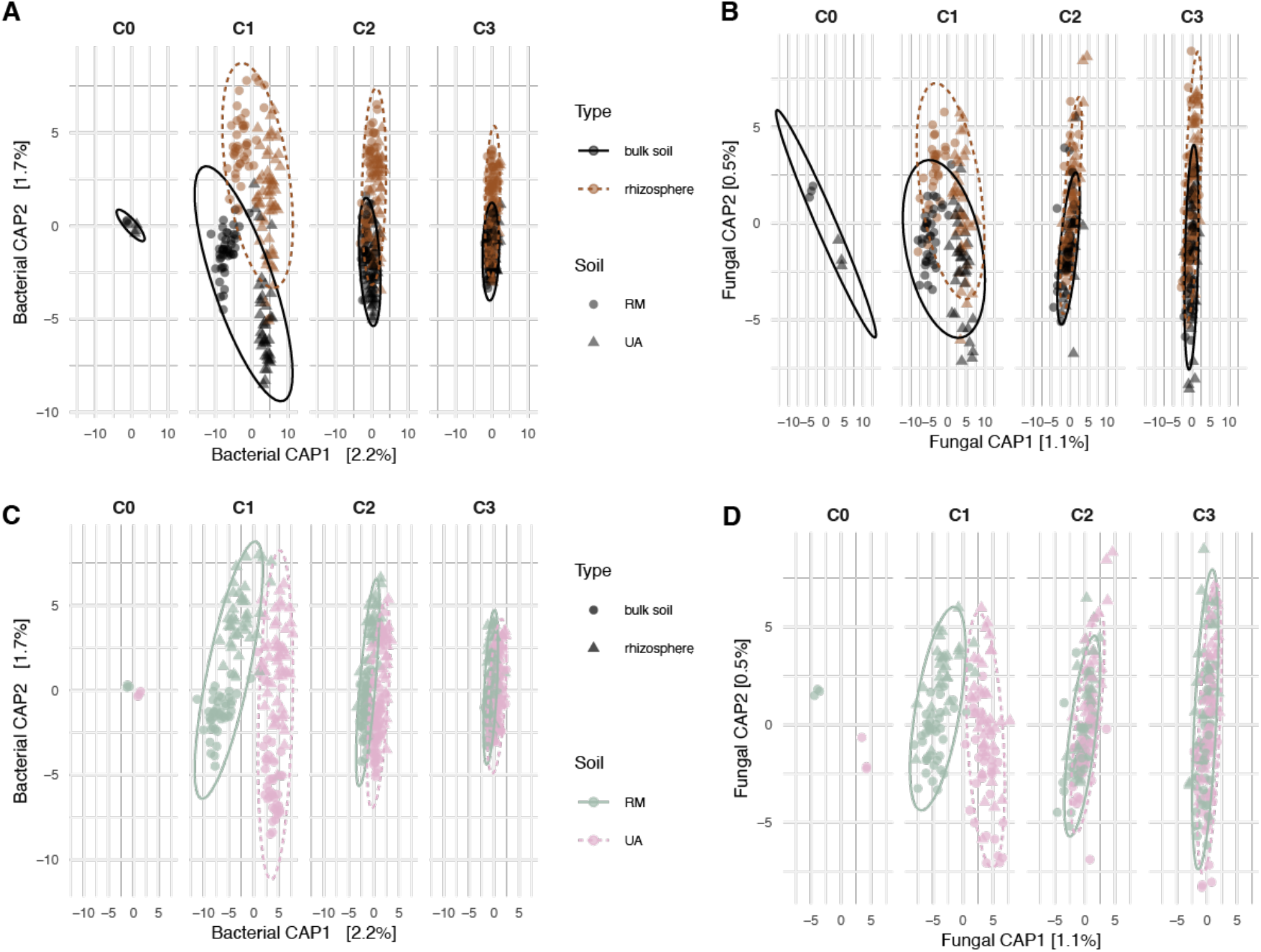
Dynamics of bacterial (left) and fungal (right) microbiomes beginning with field-collected soils (“C0”) through three cycles of passaging into sterile potting soil background (“C1-C3”). Constrained partial ordination of Aitchison distances among samples was used to remove the main effects of Cycle and sequencing depth. Constraining variables were Soil inoculum, sample type (bulk soil/unplanted *vs.* rhizosphere/unplanted), and their interaction. Points represent the taxonomic community composition, colored by (A-B) sample type and (C-D) starting soil inoculum. Ellipses denote 95% confidence intervals around the centroids.

**Figure S7.**
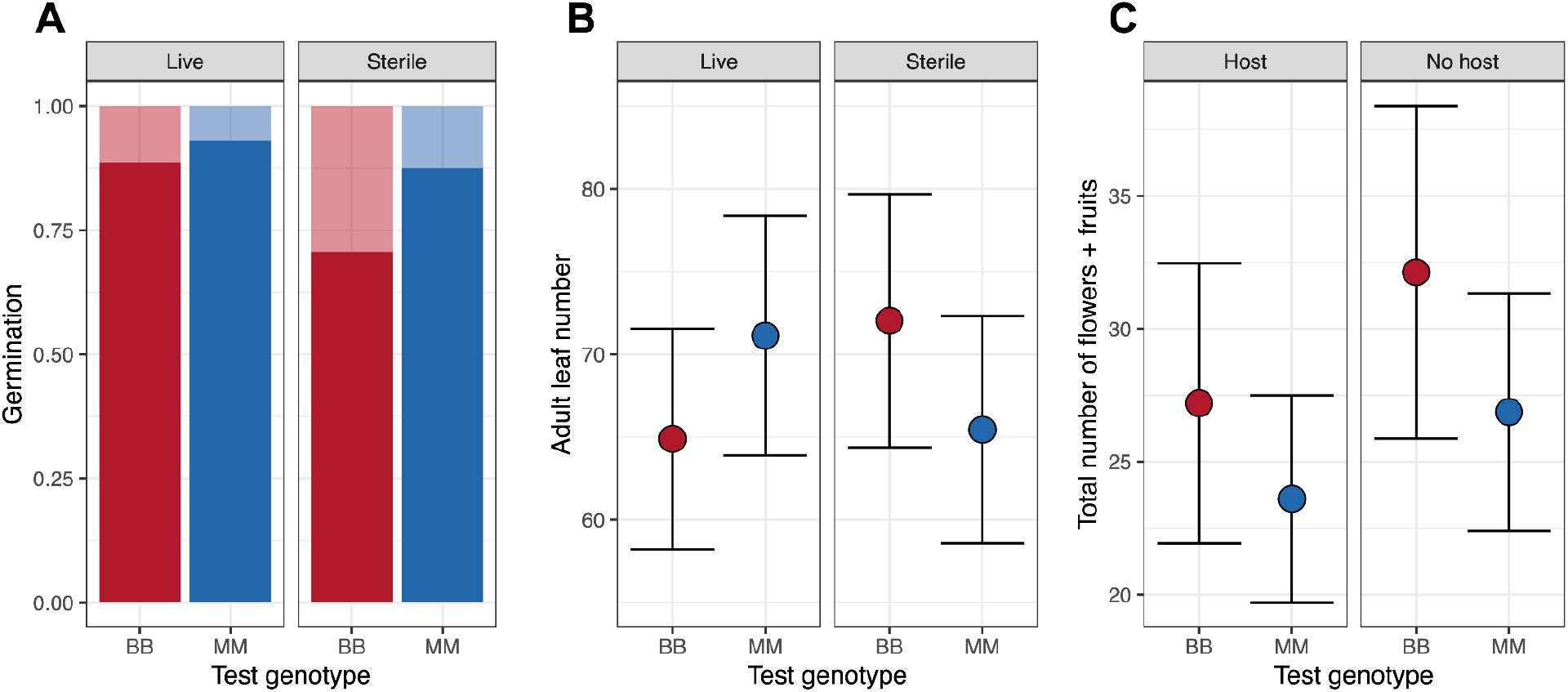
Effects of microbes writ large, agnostic to the conditioning genotype or soil source, on the expression of plant phenotypes and fitness in the test phase. **(A)** Germination proportion was marginally higher for both *BCMA* NIL genotypes when seeded into pots inoculated with live microbiomes vs. sterilized potting soil with no live inoculum added (ANOVA; main effect of microbiome: *χ*^2^=3.73, *P*=0.05341). **(B)** The expression of adult leaf number was sensitive to both plant genotype (ANOVA; main effect of genotype: *χ*^2^=7.19, *P*=0.0073) and live inoculum vs. none (main effect of microbiome: *χ*^2^=10.20, *P*=0.0014), plus their interaction (genotype*microbiome: *χ*^2^=18.15, *P*<0.0001). Because of the interaction effect, leaf number of the *BB* genotype was suppressed by live microbial inoculation relative to non-inoculated soils; patterns of expression of leaf number for the *MM* genotype were reversed. **(C)** Fecundity, estimated as the total number of flowers and fruits produced, was sensitive to host-cultivated live inoculum (ANOVA; main effect of host plant: *χ*^2^=18.22, *P*<0.0001). Both *BCMA* NIL genotypes responded the same way, producing more reproductive structures when inoculated with bulk soil microbiomes than with host-cultivated rhizosphere microbiomes. In all panes, fill color designates NIL genotype in the test phase of the reciprocal transplant experiment. In **A**, fill opacity distinguishes the proportion of pots with successful germination (opaque fill) from pots with no germination (transparent fill). In **B-C**, points are estimated marginal means from statistical models including random effects of block and family line, and error bars are +/-1SE. Full statistical support for these results is provided in table S8.

**Table S1.** Statistical analysis of bacterial and fungal microbiomes in soils conditioned by *Boechera stricta* near-isogenic lines homozygous for contrasting *BCMA* alleles. (A) *BCMA* did not alter alpha diversity of bacterial rhizosphere communities. Results are from ANOVA with Type III sums of squares. (B) *BCMA* influenced the taxonomic composition of bacterial communities both on average across the two source soils, and in a soil-specific manner. Results are from an ANOVA-like permutation test of an ordination of Aitchison distances among samples, constrained on *BCMA*, Soil, and their interaction after partialling out the effects of Cycle and Sequencing depth.

| A. Alpha diversity ($e^{\text{Shannon index}}$) | | | | | |
| --- | --- | --- | --- | --- | --- |
|  | Bacteria |  |  | Fungi |  |
| BCMA | $F_{1,272} = 1.107$ | $P = 0.29$ | | $F_{1,8} = 0.0015$ | $P = 0.97$ |
| Soil | $F_{1,272} = 1.907$ | $P = 0.17$ | | $F_{1,206} = 0.201$ | $P = 0.65$ |
| Cycle | $F_{2,273} = 28.099$ | $P = 8.02e^{-12}$ | | $F_{2,226} = 7.323$ | $P = 0.0008$ |
| Sequencing depth (logObs) | $F_{1,271} = 95.723$ | $P = 1.48e^{-19}$ | | $F_{1,227} = 2.056$ | $P = 0.15$ |
| BCMA*Soil | $F_{1,272} = 0.035$ | $P = 0.85$ | | $F_{1,205} = 7.551$ | $P = 0.0065$ |
| BCMA*Cycle | $F_{2,271} = 0.406$ | $P = 0.67$ | | $F_{2,226} = 0.716$ | $P = 0.49$ |
| Soil*Cycle | $F_{2,272} = 0.239$ | $P = 0.79$ | | $F_{2,226} = 1.612$ | $P = 0.20$ |
| BCMA*Soil*Cycle | $F_{2,271} = 0.188$ | $P = 0.83$ | | $F_{2,226} = 1.289$ | $P = 0.28$ |
| B. Taxonomic composition (Aitchison distance) |  |  |  |  |  |
|  | Bacteria |  |  | Fungi |  |
| BCMA | $F_{1,276} = 1.443$ | $P = 0.008$ | | $F_{1,233} = 0.828$ | $P = 0.86$ |
| Soil | $F_{1,276} = 5.683$ | $P = 0.001$ | | $F_{1,233} = 2.515$ | $P = 0.001$ |
| BCMA*Soil | $F_{1,276} = 1.347$ | $P = 0.013$ | | $F_{1,233} = 0.815$ | $P = 0.85$ |

**Table S2. In the soil conditioning experiment, 88 ASVs changed their relative abundances in the rhizosphere in response to host *BCMA* genotype.** Centered log ratio-transformed counts of all ASVs were fit to univariate linear mixed-effects models with the predictors *BCMA*, Soil, Cycle, and all interactions among those factors; log-transformed sequencing depth as a nuisance covariate; and experimental Block as a random intercept term. **(A)** The ASVs in this table showed evidence of sensitivity to host *BCMA* genotype (*P_FDR_*<0.05), either as a main effect or in interaction with other factors. For some ASVs, the mixed-effects model did not converge and relative abundance was re-fit using a fixed-effects model that excluded the Block term. For both fixed- and mixed-effects models, strength of statistical evidence was evaluated using ANOVA with Type III sums of squares followed by adjustment of *P-*values using the Benjamini-Hochberg false discovery rate. **(B)** For each *BCMA*-sensitive ASV, we provide the estimated marginal mean log fold change in abundance between *BB* and *MM* genotypes (logfc) and its standard error. Positive estimates indicate higher abundance in rhizospheres conditioned by *BB* genotypes. Estimates of the *BCMA* effect are provided for each of the two starting soils (“RM”, “UA”) and averaged across the two soils (“both”). All estimates are averages across the three passaging cycles. “RelAbund”: relative abundance of the ASV. “Prev”: prevalence of the ASV (proportion of samples in which the ASV was present).

**Table S3.** Summary of statistical models testing for effects of test genotype (G_T_), conditioning genotype (G_C_), starting soil environment (E_S_), and all 2- and 3-way interactions on phenotypes and fitness components in the growth chamber rhizosphere reciprocal transplant experiment. Each row reflects a separate model independently fitting the named trait or fitness component in response to all main and interactive fixed effects plus random effects of statistical block and family line (nested within *BCMA* genotype). Models of binomial data (germination) were fit using mixed effects logistic regression; models of count data (rosette leaf number, adult leaf number, number of flowers and fruits) were fit using generalized linear mixed effects regressions. All other models were fit using linear mixed effects regressions (LMER). Effects of fixed terms on response variables were tested using Type III ANOVA with Wald chi-square tests.

| Response variable<br>(N, conditional $R^2$) | Fixed effect (df) | | | | | | | | | | | | | |
| --- | --- | --- | --- | --- | --- | --- | --- | --- | --- | --- | --- | --- | --- | --- |
| | $G_T$ (1) | | $G_C$ (1) | | $E_S$ (1) | | $G_T*G_C$ (1) | | $G_T*E_S$ (1) | | $G_C*E_S$ (1) | | $G_T*G_C*E_S$ (1) | |
| | $\chi^2$ | $P$ | $\chi^2$ | $P$ | $\chi^2$ | $P$ | $\chi^2$ | $P$ | $\chi^2$ | $P$ | $\chi^2$ | $P$ | $\chi^2$ | $P$ |
| Germination<br>(161, 0.22) | 0.4474 | 0.5036 | 2.2195 | 0.1363 | 1.2862 | 0.2568 | 1.2673 | 0.2603 | 0.6129 | 0.4337 | 0.4200 | 0.5169 | 0.6285 | 0.4279 |
| Rosette leaf number<br>(140, 0.18) | 0.0210 | 0.8849 | 0.0063 | 0.9366 | 0.1972 | 0.6570 | 0.0174 | 0.8949 | 0.0009 | 0.9762 | 0.0001 | 0.9909 | 0.0505 | 0.8221 |
| Height at flowering<br>(132, 0.19) | 0.5413 | 0.4619 | 0.0714 | 0.7893 | 0.0988 | 0.7533 | 0.3543 | 0.5517 | 0.0056 | 0.9403 | 0.1200 | 0.7290 | 1.1183 | 0.2903 |
| Flowering time<br>(132, 0.13) | 0.1807 | 0.6707 | 0.4352 | 0.5094 | 0.1889 | 0.6639 | 0.2204 | 0.6387 | 0.0123 | 0.9118 | 0.2474 | 0.6189 | 1.1671 | 0.2800 |
| Adult leaf number<br>(137, 0.78) | 4.5574 | **0.0328** | 9.5404 | **0.0020** | 0.3034 | 0.5818 | 2.7833 | 0.0952 | 4.2896 | **0.0383** | 13.3977 | **0.0002** | 0.8127 | 0.3673 |
| Number of flowers + fruits<br>(161, 0.81) | 0.0033 | 0.9543 | 0.4130 | 0.5204 | 3.9637 | **0.0464** | 5.4427 | **0.0196** | 12.9412 | **0.0003** | 7.1262 | **0.0076** | 13.8421 | **0.0002** |

**Table S4.** Statistical analysis of bacterial and fungal microbiomes in the roots and rhizosphere of field-grown *Boechera stricta* near-isogenic lines with contrasting alleles at the *BCMA* locus. (A) *BCMA* altered bacterial alpha diversity in a compartment-specific manner (*BCMA*\*Type interaction, fig. S3) but had no effect on fungal alpha diversity (fig. S4). Results are from ANOVA with Type III sums of squares. **(B)** *BCMA* influenced the taxonomic composition of both bacterial and fungal communities. Results are from an ANOVA-like permutation test of a partial ordination of Aitchison distances among samples, after partialling out the effects of sequencing depth, constrained on *BCMA*, Sample Type (root *vs.* rhizosphere), Treatment (rain exclusion *vs.* control), and their interactions.

| A. Alpha diversity ($e^{\text{Shannon index}}$) | | | | | |
| --- | --- | --- | --- | --- | --- |
|  | Bacteria |  |  | Fungi |  |
| *BCMA* | $F_{1,7} = 1.21$ | $P = 0.31$ | | $F_{1,7} = 0.17$ | $P = 0.69$ |
| Type (root vs. rhizosphere) | $F_{1,111} = 107.16$ | $P = 5.77e^{-18}$ | | $F_{1,119} = 92.99$ | $P = 1.29e^{-16}$ |
| Treatment | $F_{1,82} = 0.47$ | $P = 0.49$ | | $F_{1,85} = 0.0006$ | $P = 0.98$ |
| Sequencing depth (logObs) | $F_{1,142} = 271.82$ | $P = 8.27e^{-35}$ | | $F_{1,170} = 6.23$ | $P = 0.014$ |
| *BCMA* *Type | $F_{1,86} = 5.48$ | $P = 0.022$ | | $F_{1,86} = 0.10$ | $P = 0.75$ |
| *BCMA* *Treatment | $F_{1,83} = 1.24$ | $P = 0.27$ | | $F_{1,85} = 0.53$ | $P = 0.47$ |
| Type*Treatment | $F_{1,86} = 3.58$ | $P = 0.062$ | | $F_{1,86} = 1.10$ | $P = 0.30$ |
| *BCMA* *Type*Treatment | $F_{1,86} = 0.46$ | $P = 0.50$ | | $F_{1,86} = 0.23$ | $P = 0.63$ |
| B. Taxonomic composition (Aitchison distance) |  |  |  |  |  |
|  | Bacteria |  |  | Fungi |  |
| *BCMA* | $F_{1,171} = 1.41$ | $P = 0.054$ | | $F_{1,172} = 1.45$ | $P = 0.025$ |
| Type (root vs. rhizosphere) | $F_{1,171} = 18.53$ | $P = 0.001$ | | $F_{1,172} = 17.82$ | $P = 0.001$ |
| Treatment | $F_{1,171} = 2.43$ | $P = 0.001$ | | $F_{1,172} = 2.13$ | $P = 0.002$ |
| *BCMA* *Type | $F_{1,171} = 0.91$ | $P = 0.587$ | | $F_{1,172} = 0.61$ | $P = 1.00$ |
| *BCMA* *Treatment | $F_{1,171} = 0.81$ | $P = 0.837$ | | $F_{1,172} = 1.21$ | $P = 0.11$ |
| Type*Treatment | $F_{1,171} = 1.16$ | $P = 0.169$ | | $F_{1,172} = 0.61$ | $P = 0.999$ |
| *BCMA* *Type*Treatment | $F_{1,171} = 0.71$ | $P = 0.985$ | | $F_{1,172} = 0.69$ | $P = 0.998$ |

**Table S5.**
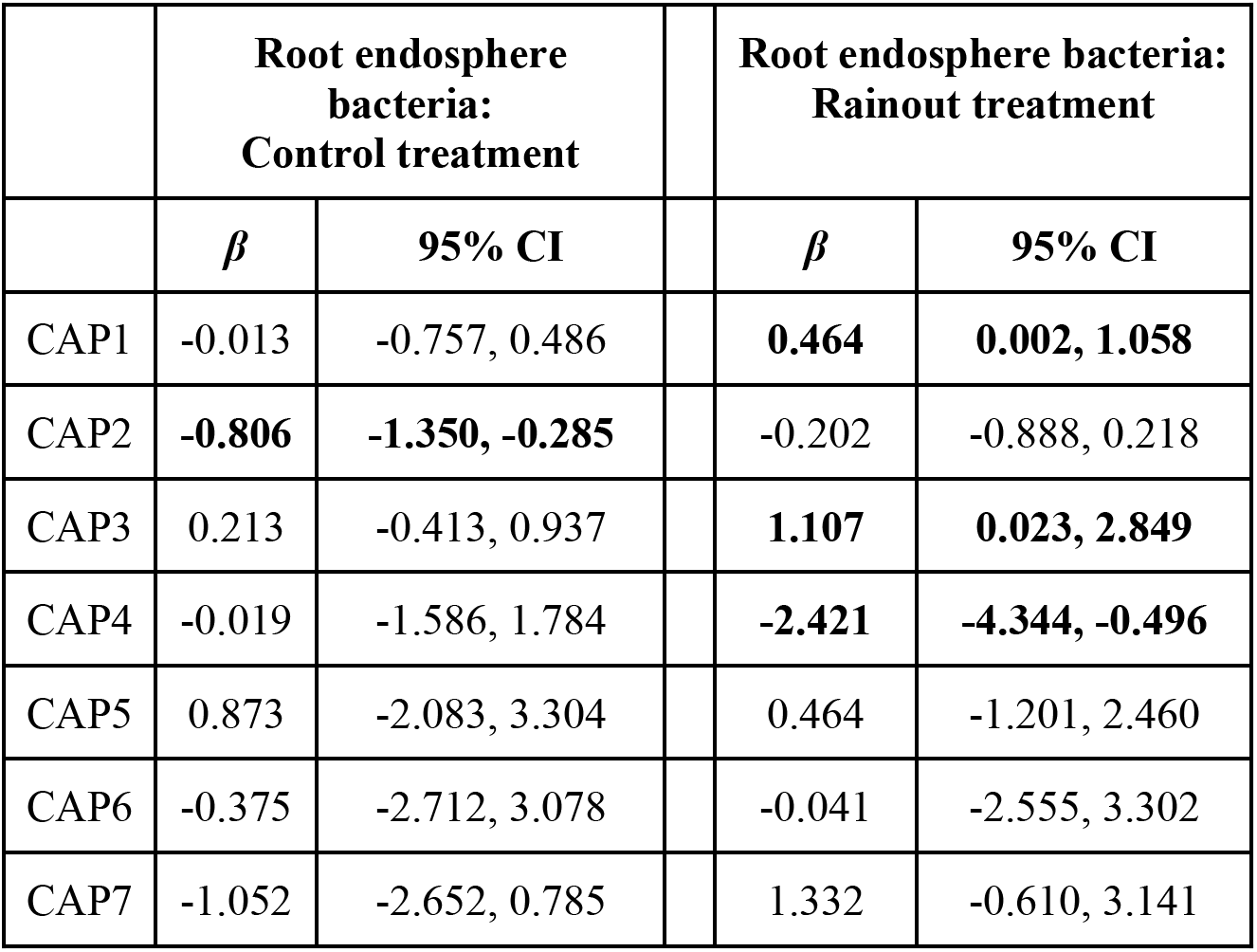
Phenotypic selection analysis on heritable components of root endosphere microbiome variation. *ꞵ* is the estimated selection gradient (the partial regression coefficient of relative lifetime fecundity) on each CAP axis in each context, derived from structural equation models. Because CAP axes were constrained on the interaction between *BCMA* genotype, treatment (control *vs.* rainout), and compartment (rhizosphere *vs.* endosphere), within each subset the CAP values describe only variation due to *BCMA* (i.e., heritable variation) and noise.

|  | Root endosphere bacteria:<br>Control treatment |  |  | Root endosphere bacteria:<br>Rainout treatment |  |
| --- | --- | --- | --- | --- | --- |
| | $\beta$ | 95% CI | | $\beta$ | 95% CI |
| CAP1 | -0.013 | -0.757, 0.486 |  | **0.464** | **0.002, 1.058** |
| CAP2 | **-0.806** | **-1.350, -0.285** |  | -0.202 | -0.888, 0.218 |
| CAP3 | 0.213 | -0.413, 0.937 |  | **1.107** | **0.023, 2.849** |
| CAP4 | -0.019 | -1.586, 1.784 |  | **-2.421** | **-4.344, -0.496** |
| CAP5 | 0.873 | -2.083, 3.304 |  | 0.464 | -1.201, 2.460 |
| CAP6 | -0.375 | -2.712, 3.078 |  | -0.041 | -2.555, 3.302 |
| CAP7 | -1.052 | -2.652, 0.785 |  | 1.332 | -0.610, 3.141 |

**Table S6.** Structural equation modeling (SEM) estimates of indirect *BCMA* effects on relative lifetime fecundity (Δ⍵) mediated by CAP axes of microbiome variation. Positive estimates of Δ⍵ indicate that selection favored the *MM* genotype over the *BB* genotype. Because CAP axes were constrained on the interaction between *BCMA* genotype, treatment (control *vs.* rainout), and compartment (rhizosphere *vs.* endosphere), within each subset the CAP values describe only variation due to *BCMA* (i.e., heritable variation) and noise.

|  | Root endosphere bacteria:<br>Control treatment |  |  | Root endosphere bacteria:<br>Rainout treatment |  |
| --- | --- | --- | --- | --- | --- |
| | $\Delta\omega_{MM-BB}$ | 95% CI | | $\Delta\omega_{MM-BB}$ | 95% CI |
| CAP1 | 0.002 | -0.168, 0.167 |  | -0.033 | -0.247, 0.111 |
| CAP2 | 0.175 | -0.153, 0.672 |  | 0.075 | -0.054, 0.524 |
| CAP3 | -0.126 | -0.631, 0.218 |  | **-0.919** | **-2.548, -0.038** |
| CAP4 | -0.011 | -0.983, 1.052 |  | **0.964** | **0.255, 2.083** |
| CAP5 | 0.118 | -0.272, 0.499 |  | -0.147 | -0.774, 0.349 |
| CAP6 | -0.078 | -0.660, 0.738 |  | 0.000 | -0.093, 0.084 |
| CAP7 | 0.011 | -0.104, 0.218 |  | 0.131 | -0.040, 0.528 |
| Total indirect | 0.089 | -1.105, 1.175 |  | 0.071 | -2.170, 1.304 |
| Total | -0.092 | -0.555, 0.405 |  | 0.129 | -0.650, 0.632 |

**Table S7.** Estimated selection gradients on bacterial ASVs’ relative abundances in the root endospheres of droughted and control plants in the field experiment.

**Table S8.**
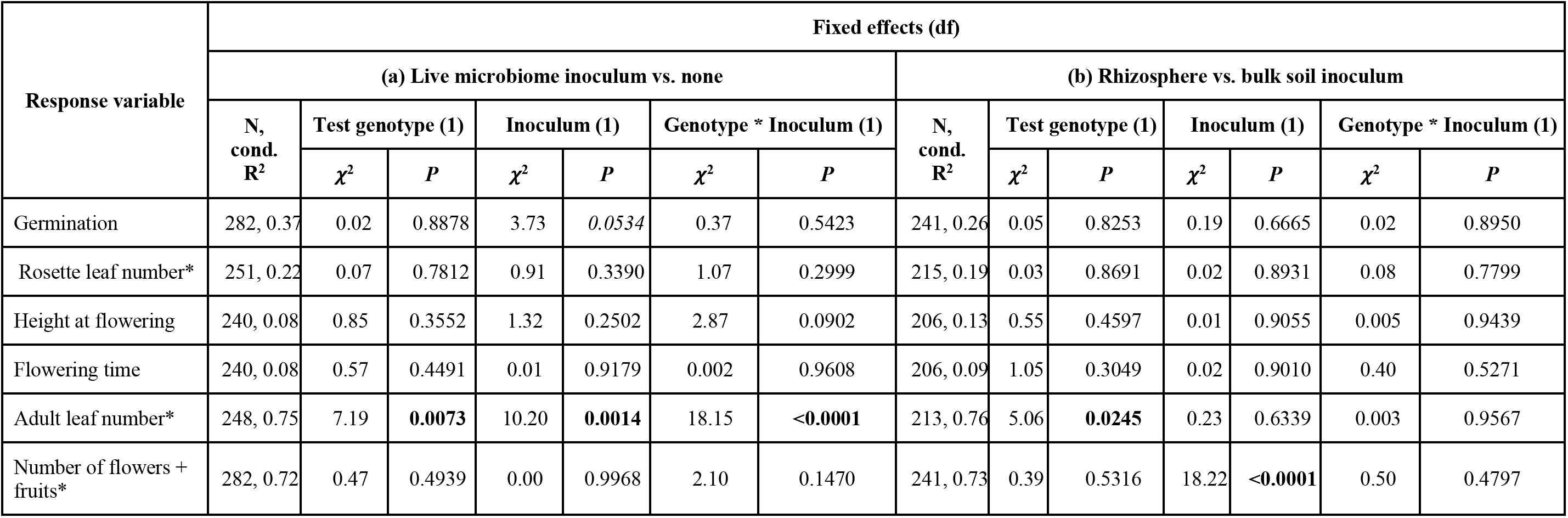
Summary of statistical models testing for effects of (a) inoculated vs. non-inoculated soils and (b) host-conditioned rhizosphere microbiomes vs. bulk soil microbiomes on genetic differences in phenotypes and fitness components in the growth chamber rhizosphere transplant experiment. Both models assess the effects of the soil microbiome broadly on plant traits in the test phase, agnostic to host genotype or starting microbial pool in the conditioning phase. Each row reflects a separate model independently fitting the named trait or fitness component in response to the main microbiome category effect plus random effects of block and NIL family line in the test phase. Models of binomial data (germination) were fit using mixed effects logistic regression; models of count data (rosette leaf number, adult leaf number, number of flowers and fruits) were fit using generalized linear mixed effects regressions. All other models were fit using linear mixed effects regressions (LMER). Effects of fixed terms on response variables were tested using Type III ANOVA with Wald chi-square tests.

| Response variable | Fixed effects (df) |  |  |  |  |  |  |  |  |  |  |  |  |  |
| --- | --- | --- | --- | --- | --- | --- | --- | --- | --- | --- | --- | --- | --- | --- |
|  | (a) Live microbiome inoculum vs. none |  |  |  |  |  |  | (b) Rhizosphere vs. bulk soil inoculum |  |  |  |  |  |  |
|  | N,<br>cond.<br>R<sup>2</sup> | Test genotype (1) |  | Inoculum (1) |  | Genotype * Inoculum (1) |  | N,<br>cond.<br>R<sup>2</sup> | Test genotype (1) |  | Inoculum (1) |  | Genotype * Inoculum (1) |  |
| | | $\chi^2$ | *P* | $\chi^2$ | *P* | $\chi^2$ | *P* | | $\chi^2$ | *P* | $\chi^2$ | *P* | $\chi^2$ | *P* |
| Germination | 282, 0.37 | 0.02 | 0.8878 | 3.73 | 0.0534 | 0.37 | 0.5423 | 241, 0.26 | 0.05 | 0.8253 | 0.19 | 0.6665 | 0.02 | 0.8950 |
| Rosette leaf number* | 251, 0.22 | 0.07 | 0.7812 | 0.91 | 0.3390 | 1.07 | 0.2999 | 215, 0.19 | 0.03 | 0.8691 | 0.02 | 0.8931 | 0.08 | 0.7799 |
| Height at flowering | 240, 0.08 | 0.85 | 0.3552 | 1.32 | 0.2502 | 2.87 | 0.0902 | 206, 0.13 | 0.55 | 0.4597 | 0.01 | 0.9055 | 0.005 | 0.9439 |
| Flowering time | 240, 0.08 | 0.57 | 0.4491 | 0.01 | 0.9179 | 0.002 | 0.9608 | 206, 0.09 | 1.05 | 0.3049 | 0.02 | 0.9010 | 0.40 | 0.5271 |
| Adult leaf number* | 248, 0.75 | 7.19 | **0.0073** | 10.20 | **0.0014** | 18.15 | **<0.0001** | 213, 0.76 | 5.06 | **0.0245** | 0.23 | 0.6339 | 0.003 | 0.9567 |
| Number of flowers + fruits* | 282, 0.72 | 0.47 | 0.4939 | 0.00 | 0.9968 | 2.10 | 0.1470 | 241, 0.73 | 0.39 | 0.5316 | 18.22 | **<0.0001** | 0.50 | 0.4797 |

